# Natural variation in the zinc-finger-encoding exon of *Prdm9* affects hybrid sterility phenotypes in mice

**DOI:** 10.1101/2023.09.06.556583

**Authors:** Khawla FN AbuAlia, Elena Damm, Kristian K Ullrich, Amisa Mukaj, Emil Parvanov, Jiri Forejt, Linda Odenthal-Hesse

## Abstract

PRDM9-mediated reproductive isolation was first described in the progeny of *Mus musculus musculus* (MUS) PWD/Ph and *Mus musculus domesticus* (DOM) C57BL/6J inbred strains. These male F_1_-hybrids fail to complete chromosome synapsis and arrest meiosis at prophase I, due to incompatibilities between the *Prdm9* gene and hybrid sterility locus *Hstx2*. We identified fourteen alleles of *Prdm9* in Exon 12, encoding the DNA-binding domain of the PRDM9 protein in outcrossed wild mouse populations from Europe, Asia, and the Middle East, eight of which are novel. The same *Prdm9* allele was found in all mice bearing introgressed *t-*haplotypes, encompassing *Prdm9* and inversions preventing recombination with wildtype Chr 17. We asked whether seven novel *Prdm9* alleles in MUS populations and the *t*-haplotype allele in one MUS and three DOM populations induce *Prdm9*-mediated reproductive isolation. The results show that only combinations of the *dom2* allele of DOM origin and the MUS *msc1* allele ensure complete infertility of intersubspecific hybrids outside the context of inbred mouse strains. The results further indicate that the erasure of PRDM9 *msc1* binding motifs may be shared by MUS mice from populations with different *Prdm9* alleles, implicating that erased PRDM9 binding motifs may be uncoupled from their corresponding PRDM9 zinc finger arrays at the population level. Our data corroborate the model of *Prdm9-mediated* hybrid sterility beyond inbred strains of mice and suggest that sterility alleles of *Prdm9* may be rare.

## Introduction

Hybrid sterility is an evolutionary concept of reproductive isolation in which hybrid zygotes develop into healthy adults that fail to produce functional gametes and are thus sterile. In the “Bateson-Dobzhansky-Muller model of incompatibilities”, hybrid sterility occurs when two or more independently evolved genes are incompatible when interacting within an individual (Bateson 1909; Dobzhansky 1936; Muller 1942). The first hybrid sterility locus identified in mammals was Hybrid sterility 1 (*Hst1*) on chromosome 17 (Forejt and Ivanyi 1974). At the *Hst1* locus, the *Prdm9* gene is responsible for the observed hybrid sterility and encodes PR domain-containing protein 9 (PRDM9) (Mihola *et al*. 2009). The PRDM9 protein is expressed in testicular tissue and fetal mouse ovaries during the early phases of meiotic prophase I when recombination is initiated (Hayashi *et al*. 2005; Lawson *et al*. 2011). PRDM9 has three conserved domains, an N-terminal KRAB domain that promotes protein-protein binding (Imai *et al*. 2017; Parvanov *et al*. 2017; Wang *et al*. 2021), an NLS/SSXRD repression domain with nuclear localization signal, and a central PR/SET domain that confers methyltransferase activity (Powers *et al*. 2016). The C-terminal domain is highly polymorphic and comprises an array of C_2_H_2_-type zinc fingers (ZNFs), which differ among PRDM9 variants in both type and number (Oliver *et al*. 2009; Baudat *et al*. 2010; Berg *et al*. 2010; Parvanov *et al*. 2010; Berg *et al*. 2011; Baudat *et al*. 2013; Buard *et al*. 2014; Kono *et al*. 2014). Variation among ZNFs is most pronounced in the amino acids at positions -1, 3, and 6 of the ZNF α-Helix, which are responsible for recognizing specific DNA target motifs (Billings *et al*. 2013; Baker *et al*. 2014; Walker *et al*. 2015; Altemose *et al*. 2017a; Altemose *et al*. 2017b; Patel *et al*. 2017). Additional amino acid substitutions in positions -5, -2, and 1 of the α-Helix are rarely seen (Parvanov *et al*. 2010; Kono *et al*. 2014). Since only amino acids at positions -1, 3, and 6 are involved in protein-DNA interactions, all other positions are not predicted to affect DNA binding affinity (Persikov and Singh 2014). Upon interaction with its specific DNA motif, the PR/SET domain of PRDM9 tri-methylates the adjacent nucleosomes on histone-3 by lysine-4 (H3K4) and lysine-36 (H3K36) (Hayashi *et al*. 2005; Wu *et al*. 2013; Eram *et al*. 2014), thereby triggering a cascade of events that initiate recombination, as reviewed in (Damm and Odenthal-Hesse 2022).

PRDM9-mediated reproductive isolation was discovered in male F_1_-hybrid progeny of *Mus musculus musculus* (MUS) and *Mus musculus domesticus* (DOM) strains that differ in *Prdm9* alleles (Forejt and Ivanyi 1974; Mihola *et al*. 2009). The PWD/Ph (hereafter PWD) MUS strain possesses the *msc1* allele, and the C57BL6/J (hereafter B6) DOM strain possesses the *dom2* allele. F_1_ hybrid males from the cross between the PWD female and the B6 male do not complete chromosome synapsis and spermatogenesis arrests at meiotic prophase I, which prevents them from forming gametes and results in a postzygotic isolation barrier (Mihola *et al*. 2009; Dzur-Gejdosova *et al*. 2012; Flachs *et al*. 2012). An X-linked locus *Hstx2*, located in a 2.7 Mb region on the proximal part of the X chromosome, modifies the effect of *Prdm9* on the fertility of intersubspecific hybrids. *Hstx2* is structurally distinct between PWD and B6 mice and causes complete hybrid sterility only when the maternal *HstX2^PWD^* is active (Bhattacharyya *et al*. 2014; Balcova *et al*. 2016). The interaction between *Prdm9* and the MUS *Hstx2^PWD^* allele promotes asynapsis of homologous chromosomes, ultimately leading to meiotic arrest (Bhattacharyya *et al*. 2013; Bhattacharyya *et al*. 2014; Balcova *et al*. 2016). In contrast, male F_1_ hybrids of the reciprocal cross (B6 × PWD) carrying the *Hstx2^B6^* allele retain a low level of fertility. The *Hstx2* locus behaves as a recombination cold spot in crosses (Balcova *et al*. 2016), has reduced *Prdm9*-mediated H3K4me3 recombination initiation sites, and lacks DMC1-decorated DNA DSB hotspots (Lustyk *et al*. 2019).

Defective pairing and high levels of chromosomal asynapsis are observed in hybrids with ineffective double-stranded break (DSB) repair (Mihola *et al*. 2009; Davies *et al*. 2016). It has been hypothesized that the molecular mechanism of PRDM9 action is related to the evolutionary divergence of homologous genomic sequences in DOM and MUS subspecies (Davies *et al*. 2016) and, more specifically, to the phenomenon of historical erosion of genomic binding sites of PRDM9 ZNF domains (Davies *et al*. 2016; Forejt 2016; Zelazowski and Cole 2016; Forejt *et al*. 2021). Nucleotide polymorphisms within genomic target motifs may affect the binding affinity of PRDM9 in heterozygous individuals. The preferential formation of DSBs occurs on the haplotype with the motif with a stronger binding affinity (Baker *et al*. 2015). However, since the uncut strand provides the template for repair, the less efficient motif is preferentially transmitted to the next generation, which can lead to the erosion of PRDM9 binding sites over time (Jeffreys and Neumann 2002). Therefore, polymorphisms that reduce the binding affinity of a given PRDM9 variant are predicted to become enriched within populations over time, resulting in the attenuation of the hotspots (Boulton *et al*. 1997). Direct evidence for over-transmission has been observed in human and mouse hotspots (Jeffreys and Neumann 2005; Berg *et al*. 2011; Cole *et al*. 2014; Odenthal-Hesse *et al*. 2014). In mouse hybrids, higher affinity binding sites for a given PRDM9 variant are four times more likely to be found on the chromosome of the other species, with which *Prdm9* did not coevolve, suggesting that binding site erosion is a predominant factor driving hotspot loss in several mouse lineages (Smagulova *et al*. 2016). In mice, about 17.5 % of hotspots have been eroded in the time it took for the PWD and B6 strains to diverge – averaging to roughly one PRDM9 binding site lost every 700 to 1500 generations (Smagulova *et al*. 2016). Indeed, in sterile hybrids of inbred strains PWD (MUS) x B6 (DOM), a bias in initiation efficiency between diverged homologs is mainly driven by functional *dom2* binding sites found on the PWD genome that are eroded on the B6 genome and vice versa for *msc1* sites (Davies *et al*. 2016), with recombination being initiated at a large number of asymmetric sets of breaks (Mihola *et al*. 2009; Davies *et al*. 2016). In this cross, fertility was restored when the B6 PRDM9 zinc-finger array was replaced with the human variant B, making symmetric recombination hotspots predominant (Davies *et al*. 2016), further supporting the hypothesis that hybrid sterility is under an oligogenic control, with PRDM9 as the main factor.

Hybrid sterility also occurs outside of laboratory models as *Prdm9* alleles found among MUS and DOM wild-derived inbred strains (Pialek *et al*. 2008) showed fertility disruption in about one-third of the intersubspecific male hybrids (Mukaj *et al*. 2020). Mice with *Prdm9* alleles that were closely related to previously identified hybrid sterility alleles showed reduced sperm counts and low paired testes weights that were associated with high asynapsis rates of homologous chromosomes in meiosis I and early meiotic arrest (Mukaj *et al*. 2020). Replacing *Prdm9^dom2^* with the ‘humanized’ targeted *Prdm9^tm1(PRDM9)wthg^ (Davies et al. 2016)* restored fertility in these mouse hybrids, supporting the role of PRDM9 as the leading player in wild-derived inbred strains (Mukaj *et al*. 2020). Furthermore, although the exon 12 sequence of *Prdm9^msc5^* had identical nucleotides in several strains, the degree of fertility reduction observed differed between strains, and the effect of the heterozygosity between genomic backgrounds remained unknown.

The relationship between the degree of chromosome asynapsis, meiotic arrest, and the number of expected symmetric DSB hotspots per chromosome was reported in PWD × B6 hybrids (Gregorova *et al*. 2018). Asynapsis was shown to operate in-*cis*, depending on the increased heterozygosity of homologs from evolutionarily divergent subspecies. Introducing at least 27 Mbs of sequence homology belonging to the same subspecies (con-subspecific homology) fully restored the synapsis of a given autosomal pair (Gregorova *et al*. 2018). *Prdm9^msc1/dom2^* also displayed a sterilizing effect in MUS x CAS hybrids, where the rate of synapsis was proportional to the level of non-recombining MUS genetic background (Valiskova *et al*. 2022).

To date, complete sterility has been observed in hybrids of inbred strains where hotspot erosion is exacerbated and the same strain-specific allele activates all hotspots (Davies *et al*. 2016; Smagulova *et al*. 2016). Through the process of inbreeding, a genome that initially possessed wildtype levels of heterozygosity becomes sequentially homozygous, and this process should equally affect PRDM9 binding sites. However, as soon as a strain is fully inbred, there should be no more heterozygous binding motifs, and erosion should cease entirely or at least slow down substantially, as it is now based only on rare mutations with stochastic placement within binding sites. Substantial erosion of binding sites must have, therefore, occurred before inbreeding, likely as a result of the high frequency of *msc1* alleles in wild populations (Buard *et al*. 2014; Kono *et al*. 2014; Forejt *et al*. 2021). In natural populations, *Prdm9* evolves rapidly, with protein variants behaving like the predator and specific motifs as prey following Red Queen dynamics to avoid negative selection of a complete loss of recombination hotspots over time (Abe *et al*. 2004; Latrille *et al*. 2017). However, while the *Prdm9* gene shows remarkable natural allelic divergence, with more than 150 alleles found in mouse populations to date (Buard *et al*. 2014; Kono *et al*. 2014; Vara *et al*. 2019; Mukaj *et al*. 2020), little is known about how many of these alleles are hybrid-sterility inducing, nor about their DNA binding motifs and their level of erosion.

Furthermore, the fertility of F_1_ hybrids could thus also be modified by additional hybrid sterility loci. At least three autosomal polymorphic hybrid sterility factors exist between PWD and STUS strains (Bhattacharyya *et al*. 2014). Five *Prdm9*-dependent quantitative trait loci have been identified in intersubspecific (MUS x CAS) hybrids, segregating on DOM background (Valiskova *et al*. 2022). However, not only do laboratory intercrosses between wild-derived inbred strains differ from the pure form of hybrid sterility observed in (PWDxB6) laboratory crosses, but contrasting patterns are also observed in wild mice. The natural hybrid zone is a relatively recent secondary contact zone across Europe, where only a third of all house mouse males exhibit fertility traits below the range of the pure subspecies. Complex polygenic control of hybrid sterility has been observed, and several interchangeable autosomal loci have been proposed to be sufficient to activate the Dobzhansky-Muller incompatibility in wild mouse hybrids (Dzur-Gejdosova *et al*. 2012; Turner and Harr 2014)*. A* genome-wide association study (GWAS) revealed strong interactions between Ch17 and Chr X, but most of these loci were located outside of *Prdm9* and *Hstx2* (Turner and Harr 2014).

Naturally occurring chromosome 17 haplotypes, the *t* haplotypes (Silver 1985) also strongly influence male fertility in wild mice. Males heterozygous for the *t*-haplotype pass it on to more than half of their offspring, with some variants presenting transmission rates over 90%, while females transmit the *t*-haplotype within the expected Mendelian ratio (Lyon 2003). Despite a strong drive, *t*-haplotypes are only present in 10-40% of all populations of wild house mice, presumably because they also include genes causing male infertility and embryonic lethality (Olds-Clarke 1997; Planchart *et al*. 2000; Schimenti *et al*. 2005; Kelemen and Vicoso 2018). Several distorter loci of the *t*-haplotype act in *trans* to impair motility in wildtype spermatozoa. They over-activate a signaling pathway controlling sperm motility kinase (SMOK), resulting in abnormal flagellar movements and loss of sperm motility (Herrmann *et al*. 1999). To prevent the distorter locus from affecting *t*-haplotype-bearing sperm, a responder locus, consisting of a fusion gene of a SMOK member and the 3’-UTR of the triple ribosomal s6 gene, confers partial cis-resistance to overactivation by the distorters, thereby restoring sperm motility (Herrmann and Bauer 2012). The *t*-haplotype consists of 30 Mb of introgressed sequence transferred from an unidentified *Mus* ancestor in the *Mus musculus* subspecies over one million years ago (Hammer and Silver 1993) and encompasses the *Prdm9* locus (TRACHTULEC *et al*. 2008), with the most diverse allele of the *Mus musculus* subspecies identified to date (Kono et al. 2014). However, it remains unknown whether *Prdm9* contributes to the observed reduction in fertility associated with *t*-haplotypes.

In summary, the low incidence of sterile wild-mouse hybrids in DOM/MUS natural hybrid zone (TURNER *et al*. 2012) together with the reported large number of hybrid sterility loci in intersubspecific backcrosses and intercrosses contrasts with the F_1_ hybrid sterility model based on the *Prdm9* allelic incompatibility, *Hstx2*, and background heterozygosity in PRDM9 binding sites. Further experimental evidence is needed to understand the mechanism of *Prdm9*-driven hybrid sterility and its role in wild mouse populations. Here, we ask whether hybrid sterility is under the PRDM9 control in wild mice beyond the context of inbred strains of mice. If the *Prdm9*-driven hybrid sterility is linked to the asymmetric erosion of PRDM9 binding motifs, then in a simple scenario, the sterility-inducing alleles would be expected to be ancestral alleles situated closest to the common ancestor on the phylogenetic tree. To test this hypothesis, we examine the evolutionary relationship between the known *Prdm9* hybrid sterility alleles and any newly identified allele.

## Material and Methods

### Mice

All work involving experimental mice was performed according to approved animal protocols and institutional guidelines of the Max Planck Society and with permits obtained from the local veterinary office ‘Veterinäramt Kreis Plön’ (permit number: 1401-144/PLÖ-004697). Mice, including strains of PWD/Ph strain, C57/Bl6 strain with transgene *Prdm9^tm1.1^(*PRDM9*)^Wthg^* strain and consomic C57BL/6J-Chr X.1s^PWD/Ph^/ForeJ mice, as well as several wild mice populations were all maintained in the mouse facilities of the Max Planck Institute for Evolutionary Biology in Plön, following FELASA guidelines and German animal welfare law. We analyzed three outcrossed populations of DOM mice; first, the French Massif-Central (MCF) population, founded in December 2005 with a starting population size of sixteen breeding pairs, with additional wild-caught animals introduced into the breeding population at the beginning of April 2010. These mice were in generation sixteen at the start of this experiment, nine generations since crossing in with the second set of new wild-caught animals. The German Cologne-Bonn (CBG) population was founded in August 2006 with ten breeding pairs and maintained as an outcross for fourteen generations at the start of this experiment. In November 2012, new wild-caught breeding pairs were crossed in. The Iranian population from Ahvaz (AHI) was started in December 2006, with six founding breeding pairs. It has been maintained in an outcross for fifteen generations at the start of this experiment, after two rounds of reduction due to inbreeding depression. Seventeen breeding pairs of mice initially trapped in Almaty, Kazakhstan (AKH) in December 2008, founded the *Mus musculus musculus* population, which had been maintained for thirteen generations at the beginning of this experiment. Whole-genome sequencing and transcriptomic data of multiple individuals from each population are publicly available in the European Nucleotide Archive and as custom tracks in the UCSC genome browser (HARR *et al*. 2016).

### Organ withdrawal

Organ withdrawal after euthanasia is not legally considered an animal experiment according to §4 of the German Animal Welfare Act. It, therefore, does not need to be approved by the competent authority (Ministerium für Landwirtschaft, ländliche Räume, Europa und Verbraucherschutz). F_1_ hybrid males were euthanized after being first rendered unconscious by deliberately introducing a specific CO_2_/O_2_ mixture ratio, then sacrificed using CO_2_ euthanasia followed by cervical dislocation. To reduce loose hair contaminating the organs during the dissection of the animal, their coat was sprayed with 75% EtOH before organ withdrawal. Spleen, a liver lobe, and both testes were extracted, and epididymides were removed. One epididymis was placed in 500 µl of cold phosphate-buffered saline for sperm counting, and all other organs were immediately snap-frozen in liquid nitrogen and stored at - 70° C.

### Fertility phenotyping

We collected three fertility parameters, body weight (BW) and paired testes weight (TW), and spermatozoa released from epididymal tissues, counted in Million/ml (SC). One epididymis, including caput, corpus, and cauda, was repeatedly cut in 1 ml of cold phosphate-buffered saline to release spermatozoa. The tube was vigorously shaken for 2 minutes, and spermatozoa in the solution were diluted to 1:40 in PBS. We counted 10 µl of diluted spermatozoa in a Bürker chamber (0,1 mm chamber height), where two replicates of 25 squares were counted. In cases when only a few (<10) spermatozoa were found, additional dilutions were prepared and counted. We added the two replicated 25 squares counts (A_25 +_ B_25_) from spermatozoa released from a single epididymis to approximate spermatozoa released from a pair of epididymides. The epidydimal spermatozoa count released in 1 ml PBS was then calculated by taking the paired counts, the volume of 25 squares (V_25_=0,02*0,02*0,01*25=0,0001 cm^3^), and the dilution factor into account.

### Spreading and Immunofluorescence analyses of spermatocytes

Spermatocyte nuclei were spread for immunohistochemistry as described in (Anderson *et al*. 1999), with the following modifications. Firstly, a single-cell suspension of spermatogenic cells from the whole testis was prepared in 0.1 M sucrose solution. The sucrose-cell slurry to which protease inhibitors (Roche 11836153001) were added, then dropped onto paraformaldehyde-treated glass slides. Glass slides were kept in a humidifying chamber for 3 hours at 4 °C to allow cells to spread and fix. Slides were briefly washed in distilled water and transferred to pure PBS before blocking in PBS with 5-vol% goat serum. Primary antibodies HORMAD2 (a gift from Attila Toth, rabbit polyclonal antibody 1:700), SYCP3 (mouse monoclonal antibody, Santa Cruz, #74569, 1:50), yH2AX (ab2893. 1:1000), and CEN (autoimmune serum, AB-Incorporated, 15-235) were used for immunolabelling. Secondary antibodies goat anti-Mouse IgG-AlexaFluor568 (MolecularProbes, A-11031), goat anti-Rabbit IgG-AlexaFluor647 (MolecularProbes, A-21245), goat anti-Human IgG-AlexaFluor647 (MolecularProbes, A-21445), goat anti-Rabbit IgG-AlexaFluor488 (MolecularProbes, A-11034) were used at 1:500 concentration at room temperature for one hour. A Nikon Eclipse 400 microscope with a motorized stage control was used for image acquisition with a Plan Fluor objective, 60x (MRH00601). Images were captured with a DS-QiMc monochrome CCD camera and the NIS-Elements program (from Nikon). Image J software was used to process the images.

### *Prdm9* Genotyping

We used ear clips taken at weaning to identify *Prdm9* allelic variation in the wild mouse populations. All F_1_ and F_2_ hybrid offspring used in the experiments were instead genotyped from the counted sperm sample taken from one of the epididymides. Furthermore, we confirmed initial parental PRDM9 genotyping after successful mating (> 5 male offspring) by sacrificing all F_0_ males. All genotyping was done on individual mouse IDs, but in such a way that the experimenter was blind to the matching fertility phenotypes.

### DNA extraction

DNA was extracted from ear clips or whole ears using salt extraction. Briefly, cells were lysed in SSC/0.2 % SDS, and proteins were digested using Proteinase K (20 mg/µl), incubating at 55 °C overnight. We salted out the DNA using 4.5 M NaCl solution, followed by two consecutive rounds of Chloroform extraction. The DNA was then Ethanol precipitated and washed twice with 70 % ethanol, and the pellet was then dried at room temperature and finally dissolved in 30 µl Tris-EDTA pH 8.0. The DNA samples were stored at 4 °C for short-term and - 70 °C long-term storage. The slurry of isolated spermatozoa with epididymal tissues was processed similarly; however, to lyse sperm heads and remove Protamines, we increased the SDS concentration to 1 % and added not only Proteinase K (20 mg/µl) but also TCEP (Thermo Scientific 77720, 0.5 M) to a final concentration of 0.01 µM. This extraction method produces a mixture of DNA extracted from somatic and sperm cells.

### Amplification of the minisatellite coding for the zinc-finger array of PRDM9

The ZNF arrays of each mouse were PCR amplified similarly as in (Buard *et al*. 2014) on 10-30 ng of genomic DNA in 12 µl reactions of the PCR buffer “AJJ” from (Jeffreys *et al*. 1990) using a two-polymerase system with Thermo Taq-Polymerase (EP0405) and Stratagene Pfu Polymerase (600159) to ensure high-fidelity PCR. When offspring are heterozygous for two alleles of different lengths (in most cases), we separated heterozygous bands after gel electrophoresis on Low Melting agarose (Thermo Fischer #R0801) by excising the bands and eluting the DNA using Agarase (Thermo Fischer #EO0461). If two heterozygous bands were apparent, excised and eluted product was immediately used in sequencing reactions after estimating the amount of DNA from the gel. If only one band was evident, alleles were not separated by size. Therefore, the purified PCR products were cloned using TOPO TA Cloning Kit for Sequencing (Life Technologies no. 450030), following the manufacturers’ specifications before sequencing. We analyzed at least eight clones per sample.

### Sequencing

Sequencing reactions of either eluted PCR product or picked clones were set up using BigDye 3.0, according to the manufacturer’s protocol, then purified using X-terminator, and finally sequenced using 3130x/ Genetic Analyzer. Only PRDM9 variants with less than 12 ZNFs could be sequenced to their ends in both directions. Exon 12 of *Prdm9* was fully sequenced for all alleles; however, forward and reverse sequences overlapped along the entire length of the exon only in alleles smaller than <1000 bp, such that larger alleles had sequence stretches only covered by either forward or reverse sequencing. Nevertheless, the sequencing products of all alleles still provided sufficient overlap for full-length assembly. We assembled the forward and reverse sequences based on the estimates of fragment sizes from PCR products on gels to accurately assemble the coding minisatellite using Geneious Software (Version 10.2-11). After sequencing and alignment, assembled minisatellites were conceptually translated into the amino acid sequence of the ZNF domain, and HMMER scores were computed using a Polynomial SVM (PERSIKOV AND SINGH 2014).

### Phylogenetic Analyses

The phylogeny on all alleles tested for hybrid sterility phenotypes in (MUKAJ *et al*. 2020) and this publication was computed using the R package “repeatR” from https://mpievolbio-it.pages.gwdg.de/repeatr/(DAMM *et al*. 2022). Briefly, minisatellite-like repeats within the gene are identified, extracted, and filtered for incomplete sequences before matrices based on minimum edit distance (Hamming) were computed using weighting costs w_mut_=1, w_indel_=3.5 and w_slippage_=1.75 as given in (VARA *et al*. 2019). These Minimum edit distances represent a metric on the set of changes between *Prdm9* minisatellite repeat units of 84 bp in length. As such, it can be used as a measure of genetic distance. We computed two distance matrices for each type of repeat, as in (DAMM *et al*. 2022). The first distance matrix included all nucleotides, while the second matrix excluded nucleotides known to be under positive selection, which are coding for the hypervariable amino acids responsible for DNA binding specificity (-1, +3, +6). Two phylogenetic reconstructions of the *Prdm9* hypervariable region were then computed separately from both matrices, using a neighbor-joining approach with the “bionj” function of the R package ape 5.0 (PARADIS AND SCHLIEP 2019) and rooted on the” humanized” *Prdm9* allele from (DAVIES *et al*. 2016).

### Genotyping for Chr17 *t*-haplotype and X-chromosomal haplotypes near *Hstx2*

The presence of the *t*-haplotype was tested using markers Tcp1 and Hpa-4ps (PLANCHART *et al*. 2000), and X-chromosomal haplotypes across the refined *Hstx2* interval were tested using primers in **Table S2** from (LUSTYK *et al*. 2019). Each forward primer was labeled with either HEX or FAM and amplified using the ABI Multiplex Kit according to the manufacturer’s protocol. Fragment lengths were then analyzed by capillary electrophoresis using a 3730 DNA Analyzer. Allele sizes were scored and binned using the Microsatellite plugin in Geneious v.10.2.

### PRDM9 *in-silico* DNA binding predictions

For *in-silico* DNA motif binding predictions, the nucleotide sequence was first conceptually translated into a protein sequence, and the C_2_H_2_ zinc-finger binding predictions were computed using a polynomial kernel with the method of (Persikov and Singh 2014). We converted the matrices from (Persikov and Singh 2014) to tab MEME, JASPAR, and STAMP input files by using the RSAT matrix conversion tool (Santana-Garcia *et al*. 2022)(http://rsat.sb-roscoff.fr/convert-matrix_form.cgi), choosing the reverse complement option. We used MEME input files for TomTom and STAMP input files for STAMP and JASPAR input files for PWMScan. The thus computed JASPAR files were inputted into PWMScan (Ambrosini *et al*. 2018) to find binding site predictions on the *Mus musculus* reference genome mm10 (which most closely resembles the genome of the C57Bl6/J strain). The .bed files containing the genome-wide putative binding sites of each PRDM9 variant were compared using bedtools intersect (Quinlan and Hall 2010), reporting each incident where bed files overlapped for at least one base pair.

### Statistical Analyses

The majority of graphs, calculations, and statistical analyses were performed using GraphPad Prism software version 9.4.1 for Mac (GraphPad Software, San Diego, CA, USA). Statistical tests are stated in the text and the Figure legends. Briefly, pairwise comparisons were performed using unpaired t-tests with Welch correction, and as we did not assume equal sample variances *a priori,* these were compared using F-tests. Similarly, multiple comparisons of fertility parameters were first evaluated for differences in sample variances using Brown-Forsythe ANOVA tests. If significant differences between means were observed, we performed Welch’s ANOVA with Dunnett’s T3 multiple comparisons test. When there was no indication of unequal sample variance, we performed ordinary one-way ANOVA instead, which we evaluated with Bonferroni multiple comparisons tests. Asynapsis data were compared between genotypes using unpaired t-tests with Welch correction, and linear correlation was assessed using the Pearson correlation coefficient (r). Using the binomial probability calculator on the VassarStats: Website for Statistical Computation (http://vassarstats.net), we tested for transmission ratio distortion.

## Results and Discussion

Previous analyses had identified four alleles of *Prdm9* that induced hybrid sterility in wild-derived inbred strains of mice initially trapped in Europe (MUKAJ et al. 2020; FOREJT et al. 2021). We screened additional European wild mice further away from the hybrid zone and mice from Asia and the Middle East for novel *Prdm9* alleles. Mice were initially caught by (HARR et al. 2016), MUS were initially trapped in Almaty, Kazakhstan (43°16’N, 76°53’E), and DOM in three different locations: the city of Ahvaz, Iran (31°19’ N, 48°42’ E), the Massif-Central area in France (45°32’N, 2°49’E) and the Cologne-Bonn area in Germany (50°52’N, 7°8’E). Previous observations of diverse haplotypes in the Iranian basin (HARDOUIN *et al*. 2015) and demographic analyses of the source populations also confirmed the AHI population as the most ancestral (FUJIWARA *et al*. 2022). These populations have been housed and maintained as outcrosses for many generations before this study (see Materials and Methods). Despite high degrees of relatedness, these populations have maintained a much larger genomic diversity than inbred strains (LAWAL *et al*. 2021). These outcrossed populations show low levels of introgression between MUS/DOM (Ullrich *et al*. 2017) and moderate levels of bidirectional introgression patterns from *Mus spretus* in all three DOM populations (Banker *et al*. 2022). These outcrossed populations of mice with inter-individual genetic diversity have high average SNP densities compared to the C57BL/6 strain, with the number of population-private variants and genomic introgression for each population collected in (**Table S1**). The distribution of original trapping locations of all mice tested for hybrid sterility phenotypes in this study and in (MUKAJ et al. 2020) are shown in **Figure S1**. Testis mRNA expression levels are available for multiple individuals of each outcrossed population (Harr *et al*. 2016) and have demonstrated a robust *Prdm9* expression (KELEMEN *et al*. 2022). In summary, these genetically diverse individuals from several outcrossed populations provide a unique resource to evaluate the *Prdm9* allelic incompatibility-mediated hybrid sterility model beyond the context of inbred strains of mice. We screened all populations for individual *Prdm9* alleles by sequencing exon12 of *Prdm9* containing the minisatellite coding for the C_2_H_2_ zinc-finger domain of PRDM9 as described in (BUARD et al. 2014; KONO et al. 2014). The amino-acid variation between ZNF domains was determined by conceptually translating the nucleotide sequence of satellite repeats into the amino-acid sequence of individual zinc fingers (**Figure S1** and **Figure S1**). We defined the ZNF variation based on the amino acids at positions −1, +3, and +6 of the alpha-helix, representing the DNA contact residues (depicted in the cartoon of **Figure 3A**). Based on variation in nucleotide repeats and the composition, order, and number of repeats, we have identified eight full-length *Prdm9* alleles in MUS mice from Kazakhstan and six in DOM from France, Germany, and Iran (shown in **Figure S1B** together with previously identified alleles from (MUKAJ *et al*. 2020)). We named these *Prdm9* alleles according to the International Committee of Standardized Genetic Nomenclature for mice (MGI) (in **Figure 4** and **Figure S1B**), registered them at <u>JAX,</u> and submitted their sequences to GenBank under Accession numbers (OQ055171-OQ055188).

We found a single, peculiar *Prdm9* allele in MUS and DOM subspecies in all four original trapping locations associated with *t*-haplotypes. In this allele, nine nucleotides are deleted in the translated amino-acid sequence of the first ZNF of the array, removing three amino acids, including one of the zinc-binding Cysteine ligands (**Figure S1**). In addition, distinct amino acids are seen in positions −1, +3, and +6, not present in any other PRDM9 variants, such as TDK and ASQ, with additional differences between the two types of ASQ ZNFs in positions -8 and +5. Similarly, the ANQ ZNF found in mice with *t*-haplotypes differs from the ANQ found in mice without *t*-haplotypes at position -2, a pattern previously seen in mice with *t*-haplotypes (KONO *et al*. 2014). We, therefore, tested additional mice with *t-*haplotypes for *Prdm9*, including *Mus musculus castaneous* (CAS) from Taiwan and mouse strain *T/t^p4^* (FOREJT et al. 1988), confirming that all possessed the same *Prdm9* allele.

Given that a single *Prdm9* allele was found in all mice carrying a *t*-haplotype, regardless of subspecies, we consider it an intraspecies *Mus musculus t*-haplotype *Prdm9* allele 1 “*Prdm9^mmt1^*”. Together with *t-*haplotypes, the *mmt1* allele was always present in a heterozygous state and occurred in all of our outcrossed populations at high frequencies. We found a *t*-haplotype in 50% of the MUS population from Almaty (Kazakhstan), in 88% of the DOM population from Ahvaz (Iran), in 90% of the mice from Cologne-Bonn (Germany), and 100% of the mice from the Massif-central (France) population. Even though initial population frequencies may have gotten heavily distorted due to the *t*-haplotype over-transmission, and the alleles may not reflect the initial population frequencies in which they occurred in the wild. However, the alleles we identified in the outcrossed population should still reflect the *Prdm9* alleles in the wild. While most alleles are novel, the *mmt1* allele (KONO et al. 2014) and three MUS alleles which we found in mice from Kazakhstan had been identified previously, such as *msc6*, previously named Ma8 and identified in Grozny, Russia (KONO et al. 2014)*, msc12* allele identified as 7mus1 in MUS from (BUARD et al. 2014), and *msc9* found in strains CHD and BLG2, named Ma12 (KONO et al. 2014). A single DOM allele has been previously identified as 16dom1 (BUARD et al. 2014) in the DOT strain, originally from Tahiti (French Polynesia), which we found in Ahvaz, Iran, as *dom8*. All alleles in this study, except the *mmt1* allele, are present only in one population, yet some closely resemble alleles of other subspecies. For example, *msc11* closely resembled two alleles from the CAS subspecies, firstly, the classical *cst1* allele, which possesses Serine at position -1 of the alpha-helix of the 8^th^ ZNF (PARVANOV et al. 2010); in this position*, msc11* has a single amino-acid change to Asparagine, and secondly, a CAS trapped in Nowshahr, Iran, Ca1 (KONO et al. 2014), differs only by a single amino-acid substitution in position 6 of the alpha-helix of the 6^th^ ZNF, where Ca1 possesses Glutamine, instead of Lysine. Alleles *msc7* and *msc10* appear similar and share similarities to MUS 27mus1 trapped in Bulgaria (BUARD *et al*. 2014) and CAS Cc4 trapped in Grozny, Russia (KONO et al. 2014). The observations of CAS-like alleles in MUS populations are consistent with observations of many “MUS-like CAS” and “CAS-like MUS” samples in genomic datasets that show admixture patterns (FUJIWARA *et al*. 2022).

### Testing *Prdm9* alleles for sterility phenotypes in intersubspecific hybrids

To investigate whether newly identified wild *Prdm9* alleles induce hybrid sterility in intersubspecific crosses, we adapted the crosses used in the laboratory models of hybrid sterility to eliminate possible variation due to the *Hstx2* modifier. PWD (MUS) females were crossed with wild DOM males to test DOM alleles, emulating the PWD × B6 laboratory model. To test MUS alleles, we emulated the B6.DX1s × PWD laboratory model (MUKAJ *et al*. 2020) and crossed wild MUS males to C57BL/6J-Chr X.1s^PWD/Ph^/ForeJ females (abbreviated B6.DX1s). B6.DX1s is a consomic DOM strain of C57Bl6/J background that carries the *Hstx2* locus within a 69.6 Mb PWD sequence of the proximal end of the X-chromosome, essential for F_1_-hybrid sterility (BHATTACHARYYA *et al*. 2014; BALCOVA *et al*. 2016; LUSTYK *et al*. 2019; FOREJT *et al*. 2021).

All nine DOM sires and five of the sixteen MUS sires possessed *t*-haplotypes. The *t-* haplotype is a known meiotic driver, skewing transmission against wildtype Chr17 in the male germline. We, therefore, experienced a severe reduction of testable *Prdm9* alleles on wildtype Chr17 in both MUS and DOM mice because of the over-transmission of the *Prdm9^mmt1^*allele within the *t*-haplotype. Indeed, more than 87% of offspring from fathers with *t*-haplotypes inherited the *mmt1* allele, a significant deviation from the Mendelian 50:50 transmission ratio (two-tailed binomial probability *P=* <0.000001, approximated via normal). In contrast, all other *Prdm9* alleles were transmitted at 50:50 in mice without *t*-haplotypes. To estimate the effect of *Prdm9* on the fertility of hybrids, we used the weight of paired testes and the number of spermatozoa in paired epididymis as a proxy. Since these fertility parameters differ with the genetic background (WIDMAYER *et al*. 2020), we first looked into the physiological variation of the outcrossed source populations of wild mice as a control, which we compared with inbred mice (**Figure S1**). Wild mice from all three DOM populations had comparable sperm counts, while testes weights of CBG and MCF populations were significantly higher than in B6 mice. Similarly, wild MUS had elevated testes weights compared to PWD. Outcrossed mice also possess large variability in body weight, and we noticed that testes weight correlated with sperm count more robustly than testis weight normalized to animal body weight (**Figure S1**). Because wild mouse sires came from outcrossed populations, they were almost always heterozygous for different *Prdm9* alleles. Several males from the same population shared one of the *Prdm9* alleles. As a result, offspring from different crosses with such fathers acquired the same combination of *Prdm9* alleles. We used them to determine whether differing genetic backgrounds affect fertility parameters in F_1_ hybrid offspring with the same *Prdm9* allelic combination inherited from different fathers. We did not observe a significant background effect on sperm counts (**Figure S1A)**, except in offspring that inherited *t*-haplotypes (**Figure S1B**). Paired testes weights, however, were significantly different in MUS males with *msc6* and *msc10* alleles and hybrid males carrying *t*-haplotypes (**Figure S1**). We analyzed the fertility parameters for all mice aged 60-100 (±2) days, as hybrid fertility parameters can vary with age, with a marked decline after 20 weeks (WIDMAYER *et al*. 2020). We performed regression analyses of age and fertility for each *Prdm9* allele separately and for all alleles combined and detected no apparent effect of age on fertility in the tested age ranges, with only a weak positive correlation of age on testis weights in offspring with the *mmt1^KH^ and msc10* alleles (**Figure S1**). When we pooled fertility parameters of F_1_-hybrid males by *Prdm9* genotype, almost all had significantly higher TW and significantly elevated SC compared to control hybrid sterility crosses PWD × B6 and B6.DX1s x PWD. Exceptions are intersubspecific hybrids of PWD x DOM crosses that inherited the paternal *mmt1^MC^* allele and intersubspecific hybrids of B6.DX1s x MUS crosses that inherited the paternal *msc11* and *msc12* alleles (**Table S *3*** and **Table S *4*)**, whose SC was not significantly elevated compared to *msc1*. The hybrids inheriting paternal *msc12* showed significantly reduced fertility compared to other tested alleles (**Figure 1**). Given that all F_1_-hybrid males were either completely fertile or showed only reduced fertility, we wanted to test whether they would display chromosomal asynapsis, a hallmark characteristic of *Prdm9*-dependent hybrid sterility (FOREJT AND JANSA 2023). We performed immunofluorescence analyses on spermatocyte spreads of F_1_-hybrids that inherited alleles *msc10*, *msc11*, and *msc12* and hybrids that inherited the *mmt1* allele in combination with both DOM and MUS *t*-haplotypes.

**Figure 1.**
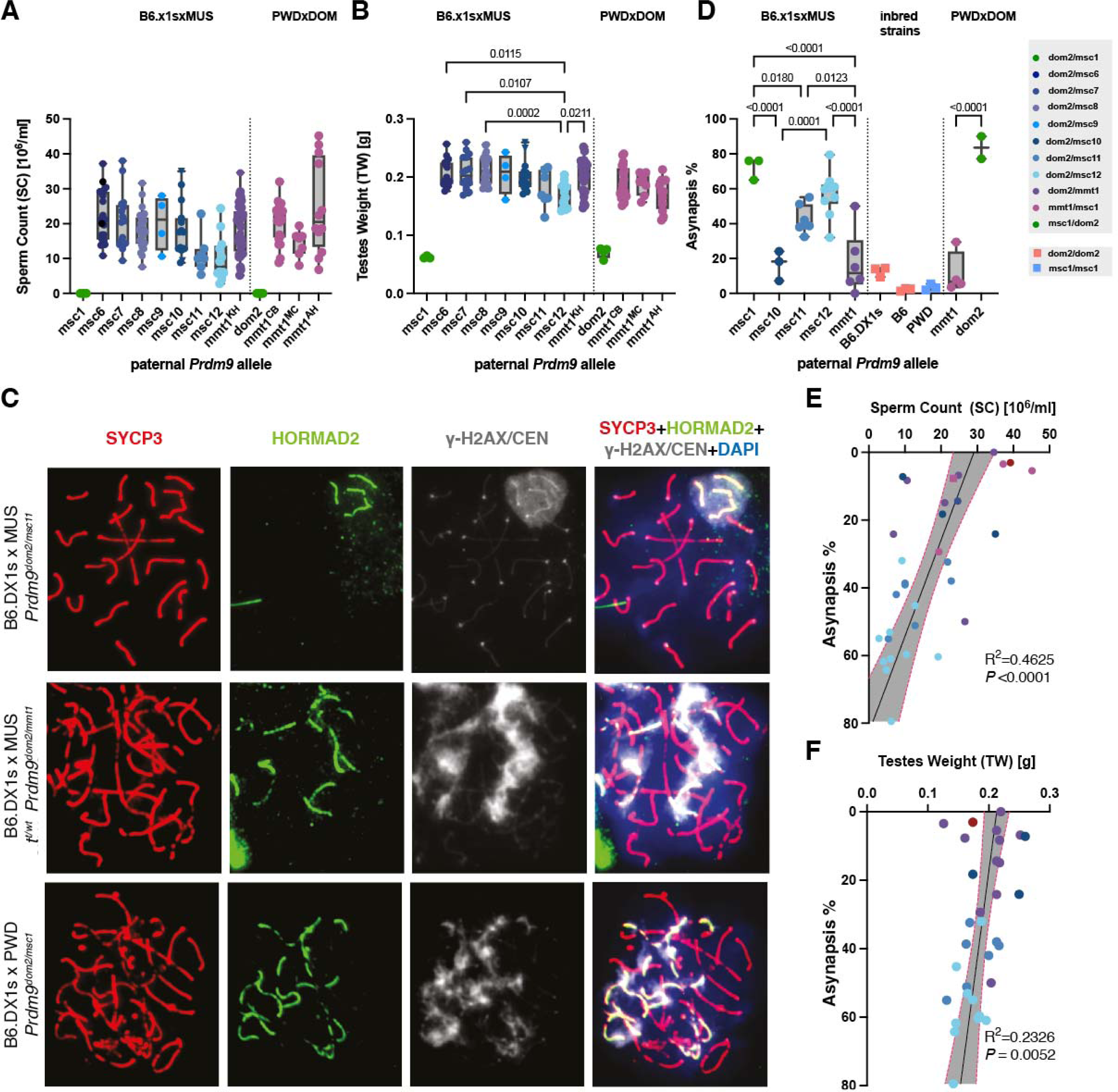
Fertility parameters of intersubspecific hybrids were grouped by *Prdm9* genotype. **(A)** Sperm count, and **(B)** paired testes weights for intersubspecific offspring of (B6.DX1s × wild MUS) or (PWD × wild DOM) crosses grouped by *Prdm9* genotype and compared to offspring of known hybrid sterility crosses (B6.DX1s × PWD) and (PWD × B6), using pairwise ANOVA with Bonferroni correction. All hybrid males carry the *Hstx1^PWD^* allele on Chr X. **(C)** The panels show spermatocyte spreads of two intersubspecific B6.DX1s × wild MUS hybrids, with differing *Prdm9* genotypes. The defects in chromosome asynapsis were assessed by antibody staining for HORMAD2 protein (green), which marks asynapsed autosomal chromosomes in addition to the nonhomologous parts of XY sex chromosomes that are physiologically observed in normally progressing meiocytes. DNA is counterstained with DAPI (blue). Synaptonemal complex assembly was evaluated by SYCP3 protein immunostaining (red) and the presence of yH2AX (grey). At the zygotene/pachytene transition, clouds of yH2AX mark chromatin associated with asynapsed axes. In addition, localized grey dots represent CEN-labeled centromeres. **(D)** The percentage of asynaptic cells on the Y-axis were grouped by *Prdm9* genotype. The percentage of asynaptic cells correlated with fertility parameters of intersubspecific F_1_-hybrids, namely **(E)** sperm count and **(F)** paired testes weights, with red dotted lines representing 95% confidence intervals, and ***P*** values, and Pearson values **r**, given on the bottom right.

To determine if cells are in the pachytene stage when the synaptonemal complex is fully formed, we immunostained synaptonemal-complex protein 3 (SYCP3). In contrast to continuous SYCP3 staining in pachytene, SYCP3 staining is disorganized and patchy in the preceding late zygotene stage (when the synaptonemal complex is still forming) and the succeeding early diplotene stage (when the synaptonemal complex begins to disassemble as autosomes de-synapse and lateral elements separate), when the SYCP3 signal becoming visible as pair of thinner threads (DE LA FUENTE *et al*. 2007). Additionally, we assessed H2AX, which localizes to the X and Y chromosomes in males only at mid-to-late pachytene, forming a punctate “sex body” of transcriptional silencing (FERNANDEZ-CAPETILLO *et al*. 2003). We evaluated 48-113 of such pachynemas for asynapsis in each individual, scoring each HORMAD2 stained element (excluding sex chromosomes) as one asynapsis event. We determined the percentage of cells with asynapsis (and collected all data in the linked <u>Dryad Repository</u>). The F_1_ hybrid males with wild MUS alleles *msc11* and *msc12* had an elevated proportion of asynaptic pachynemas (*msc11,* 42.4 ± 8.0 % and *msc12* 57.2 ± 12.5%), these frequencies are significantly elevated, compared to control mice of pure parental subspecies (B6.DX1s and PWD), with asynapsis levels of *msc12* statistically indistinguishable from those of *msc1* (**Figure 1C**). In contrast, hybrids with wild mouse allele *msc10* had lower asynapsis, averaging 16.6 ± 8.6% of cells with asynapsis, not significantly different from fertile controls and significantly lower than sterile controls (**Figure 1C**). Similarly, mice with *t-*haplotypes also showed low asynapsis rates (DOM *mmt1* 11.5 ± 12%, MUS *mmt1* 17.4 ± 18.0%). We also tested the relationship between fertility parameters correlated and the meiotic asynapsis rate across all genotypes. We observed a weak but significant correlation between high asynapsis with both low sperm count (Pearson R^2^ = 0.46, *P <* 0.0001) (**Figure 1E**) and paired testes weights (Pearson R^2^ = 0.23, *P=* 0.0052) (**Figure 1F**). In summary, the tested *Prdm9* allelic combinations were either completely fertile or showed a reduction of fertility. As hybrids with *msc11* and *msc12* alleles had the lowest sperm counts and paired testes weights and also showed the highest levels of chromosomal asynapsis, we can conclude that *Prdm9* likely drives this effect, but any *Prdm9*-dependent effect is either not binomial, affected by overall genomic heterogeneity, or by potential genetic modifiers on the wild genetic backgrounds.

### The relationship between *Prdm9* genotype and wild-derived outbred genetic background

According to the hypothesis linking the *Prdm9*-driven hybrid sterility to the asymmetric erosion of PRDM9 binding motifs, the sterility of F_1_ hybrids results from the erosion of MUS PRDM9^msc1^ binding sites in the PWD genome and DOM PRDM9^dom2^ sites in the B6 genome (Davis et al. 2016). If a given MUS population with an unrelated “fertility” *Prdm9* allele carries the same *msc1* binding motif as PWD mice, then the fertility of (PWD x MUS) x B6 hybrids would segregate according to *Prdm9*; *Prdm9^msc1^* males would be sterile, and their *Prdm9^MUS^*siblings would be fertile. However, if the given wild MUS genome does not carry any PRDM9 binding site erasure or the erased motifs match an unrelated *Prdm9* allele, then *Prdm9^msc1^*and *Prdm9^MUS^* male progeny should be fertile. The same assumption can be made for the outcome of PWD x (B6 x DOM) testcross to test the coupling of eroded PRDM9 binding motif with corresponding PRDM9 allelic zinc finger domains. To test the decoupling of eroded PRDM9 binding sites from the corresponding allelic form of *Prdm9* in wild populations, we generated intraspecific (DOM x B6) hybrid males and crossed them to PWD females to test the DOM-derived *mmt1* and *dom2* alleles against the mixed DOM background. To test MUS alleles and their genetic background, the B6.DX1 females were crossed with intraspecific (PWD × MUS) F_1_ hybrid males. We also performed an analogous cross in reciprocal orientation for MUS alleles, where (PWD × wild MUS) F_1_ hybrid females were crossed with B6 males. In the female germline, the *mmt1* allele showed even transmission (two-tailed exact binomial probability, *P =* 0.136), confirming male-specific *t*-haplotype transmission distortion (LYON 2003). Since not only *Prdm9* but also X-chromosomes segregate in this cross, we only included males with the PWD haplotype containing the refined *Hstx2*^PWD^ locus, as defined by alleles of the X chromosomal microsatellite markers SR51, SX69084, and SX65100 (LUSTYK *et al*. 2019)(see also **Figure S1**). Siblings that inherited the *Prdm9* alleles *msc6, msc7, msc8,* or the *mmt1* of MUS or DOM all displayed fertility phenotypes within the physiologically normal range. In contrast, their brothers with the allelic combination *msc1/dom2* were sterile (**Figure 2**), with testes’ weight and sperm count comparable to (PWD x B6) F_1_ males with the same allelic combination. These results suggest that mice from MUS populations with different *Prdm9* alleles may share the same pattern of erased PRDM9^msc1^ (PWD) binding motifs, implicating uncoupling of the erased PRDM9 binding motifs and PRDM9 zinc finger arrays at the population level. Furthermore, the partially wild-derived outbred background does not appear to carry additional major genetic modifiers that would prevent sterility *per-se*.

**Figure 2.**
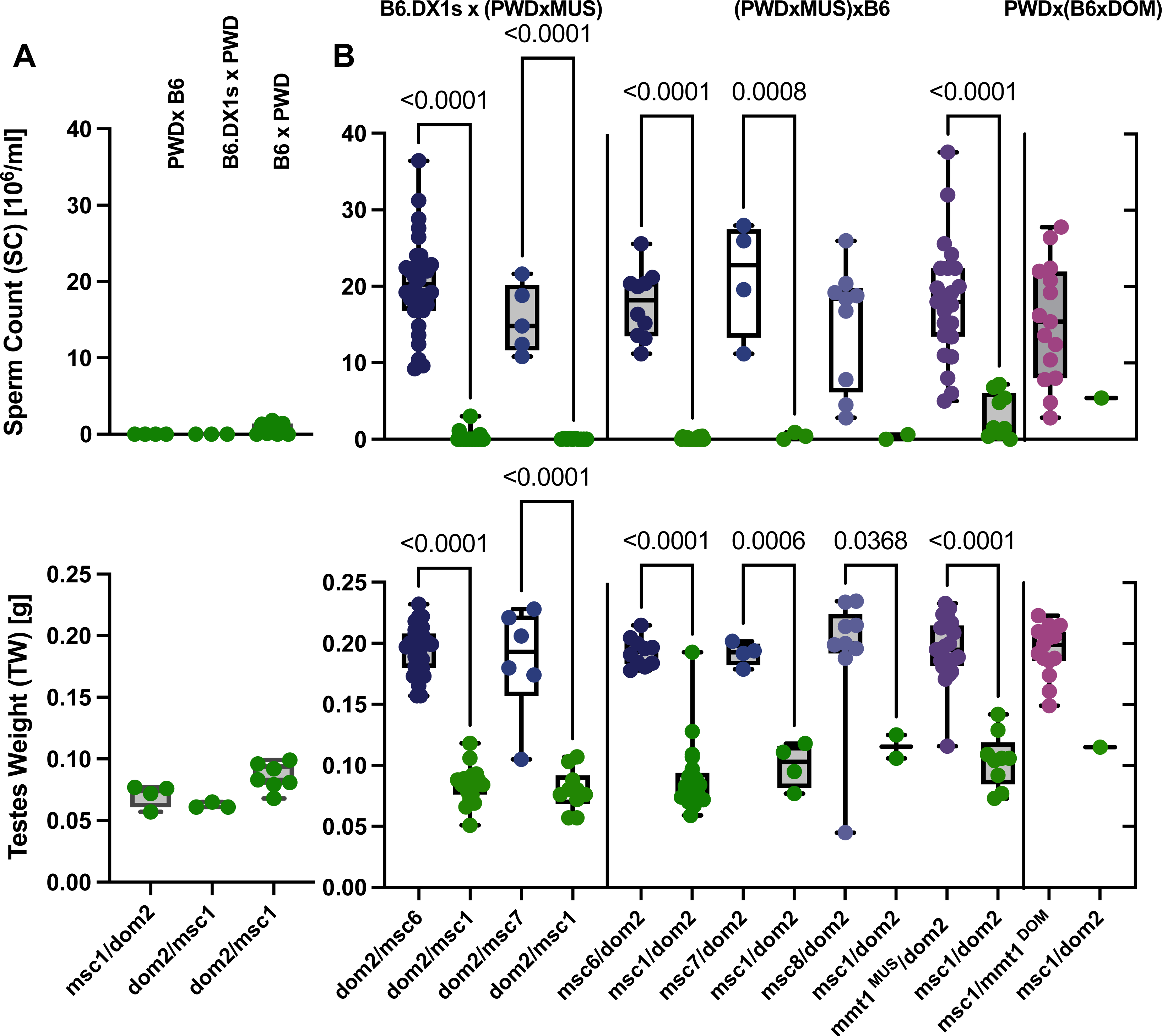
Fertility phenotypes segregate with parental *Prdm9* alleles in reciprocal intersubspecific hybrids. **(A)** Fertility parameters of control cross with the *Prdm9* allelic combination *dom2/msc1* and Hstx2 allele from PWD or B6. (B) Intersubspecific F_1_ male offspring of **(left)** B6.DX1s females crossed to intrasubspecific MUS males **(middle)**, Intraspecific MUS hybrid females crossed to B6 males **(right)** PWD females crossed to intrasubspecific DOM males. Data were pooled from parents with the same *Prdm9* genotype. Color and statistics, as in Figure 1.

### Variation of X-chromosomal haplotypes in *Prdm9-mediated* sterility

Until now, information on the possible intrasubspecific variation of the Hstx2 locus has been lacking. Since the interval 65-69 Mb on Chr X containing the Hstx2 locus appears as a recombination cold spot (BRICK et al. 2018; LUSTYK et al. 2019), we focused on this region in the Kazakh (KH) MUS population and identified four different MUS X-chromosomal haplotypes Kha, KHb, KHc, and KHd using only three microsatellite markers (see **Figure S1**) indicating that recombination events within the Hstx2 locus could have occurred. Next, we tested whether these wild X-chromosomal haplotypes differed in the modulation of Prdm9-driven hybrid sterility in crosses where Prdm9 and Hstx2 segregated. As previously observed with the Hstx2^PWD^ sterility allele, fertility co-segregated predominantly with the Prdm9 genotype, irrespective of which X-chromosomal haplotype the offspring possessed (**Figure S1A-C**). Regardless of these X-chromosomal haplotypes, all F_1_-hybrids inheriting any wild MUS Prdm9 alleles from their mothers were fertile, whereas siblings inheriting the msc1 alleles were sterile. To conclude, the Hstx2 did not show functionally defined intraspecific polymorphism within the studied MUS population (**Figure S1D**) and the results did not reveal other genetic modifiers on any individuals’ wild genetic background along the maternal germline. An exception to this rule was the Prdm9^msc1/dom2^ male offspring, whose mothers were t-haplotype carriers, which produced an average of 2.7 M spermatozoa. These males differed significantly in both sperm count and testes weight from mice with identical genotypes whose mothers did not carry t-haplotypes (Mann-Whitney test; SC P < 0.0001, TW P = 0.0094) (**Figure S1E**), suggesting the presence of Prdm9 fertility modifier(s) in t-carrying populations. The apparent trans-effect of Prdm9, located on the t-haplotype, cannot be disentangled from that of other fertility modifiers within a single haplotype block. However, while testis Prdm9 expression levels in mice with t-haplotypes were not significantly different from those in mice without t-haplotypes, several other genes on t-haplotypes are enriched for copy gain events (KELEMEN AND VICOSO 2018) and show overexpression in the testis of t^t/wt^ heterozygous mice (KELEMEN et al. 2022). However, since the *msc1/dom2* allelic combination leads to sterility even on variable genomic backgrounds, we can conclude that hybrid sterility is indeed under oligogenic control, with *Prdm9* as the leading player.

### Phylogenetic analyses of *Prdm9* alleles in mice

It has been proposed that the role of *Prdm9* in hybrid sterility could be related to the evolutionary divergence of homologous genomic sequences in DOM and MUS subspecies (DAVIES *et al*. 2016) and, more specifically, to the phenomenon of historical erosion of genomic binding sites of PRDM9 ZNF domains (BAKER *et al*. 2015; SMAGULOVA *et al*. 2016) caused by repeated biased gene conversion. Consequently, only the *Prdm9* alleles that have been present for longer evolutionary timescales should generate such partial erosion of their ZNFs binding motifs. To enquire into the evolutionary history of *Prdm9* alleles, we analyzed the phylogenetic relationship of alleles present in our wild mice populations and other alleles for which *Prdm9*-mediated hybrid sterility had been studied (PARVANOV *et al*. 2010; MUKAJ *et al*. 2020). As an outgroup, the humanized *Prdm9* “B-allele” was added (DAVIES *et al*. 2016). Since handling sequence repeats is challenging for multiple-sequence alignment algorithms, particularly when the number of repeat units differs, the allelic divergence of minisatellite sequences could not be assessed by standard assembly programs. In addition, genetic distance models (i.e., Tamura-Nei (TAMURA AND NEI 1993)) do not accurately reflect minisatellite evolution driven by *de-novo* recombination between repeats (JEFFREYS *et al*. 2013). Therefore, to reflect *Prdm9* evolution more accurately, we applied an algorithm that computes Hamming Distances between minisatellite repeat, which takes not only point mutations and small indels but also within-repeat-unit processes (w_mut_=1), as well as repeat-unit insertions and deletions (w_indel_=3.5) and even repeat-unit duplication and slippage (w_slippage_=1.75) into account (VARA *et al*. 2019; DAMM *et al*. 2022). For a more conservative phylogenetic analysis, we only included nucleotide repeats that, when translated into amino acids, had a Hidden Markov Model (HMMER) bit score above 17.7, determined using (PERSIKOV AND SINGH 2014). This removed the nucleotide repeat coding for the first zinc finger in the ZNF array, which we found to be conserved in all *Prdm9* alleles except *mmt1*.

In the neighbor-joining tree of Hamming distances rooted on the “humanized” *Prdm9* allele, alleles mostly cluster according to mouse subspecies (as shown in **Figure 3**). However, not all alleles follow the MUS/DOM subspecies divide. The *mmt1 Prdm9* allele found in all mice with *t-*haplotypes formed a separate branch irrespective of subspecies and mouse origin, a pattern typical of introgression. The large degree of conservation of *Prdm9* on the *t*-haplotype stands in stark contrast to the rapid evolution of *Prdm9* alleles in mice. Given the remarkable divergence of *Prdm9* alleles in all natural populations studied to date (BUARD *et al*. 2014; KONO *et al*. 2014; VARA *et al*. 2019; MUKAJ *et al*. 2020), a single ancestral introgression event of *t*-haplotypes into an antecedent of all *Mus musculus* subspecies appears more likely than repeated introgression of the same allele at multiple independent events. Furthermore, the frozen pattern of zinc fingers in PRDM9^mmt^ can be explained by a series of naturally occurring inversion blocks, one including the *Prdm9* locus (KELEMEN AND VICOSO 2018), likely causing recombination suppression in t-haplotypes, therefore constraining *Prdm9* evolution that is mainly recombination-driven (JEFFREYS *et al*. 2013).

**Figure 3.**
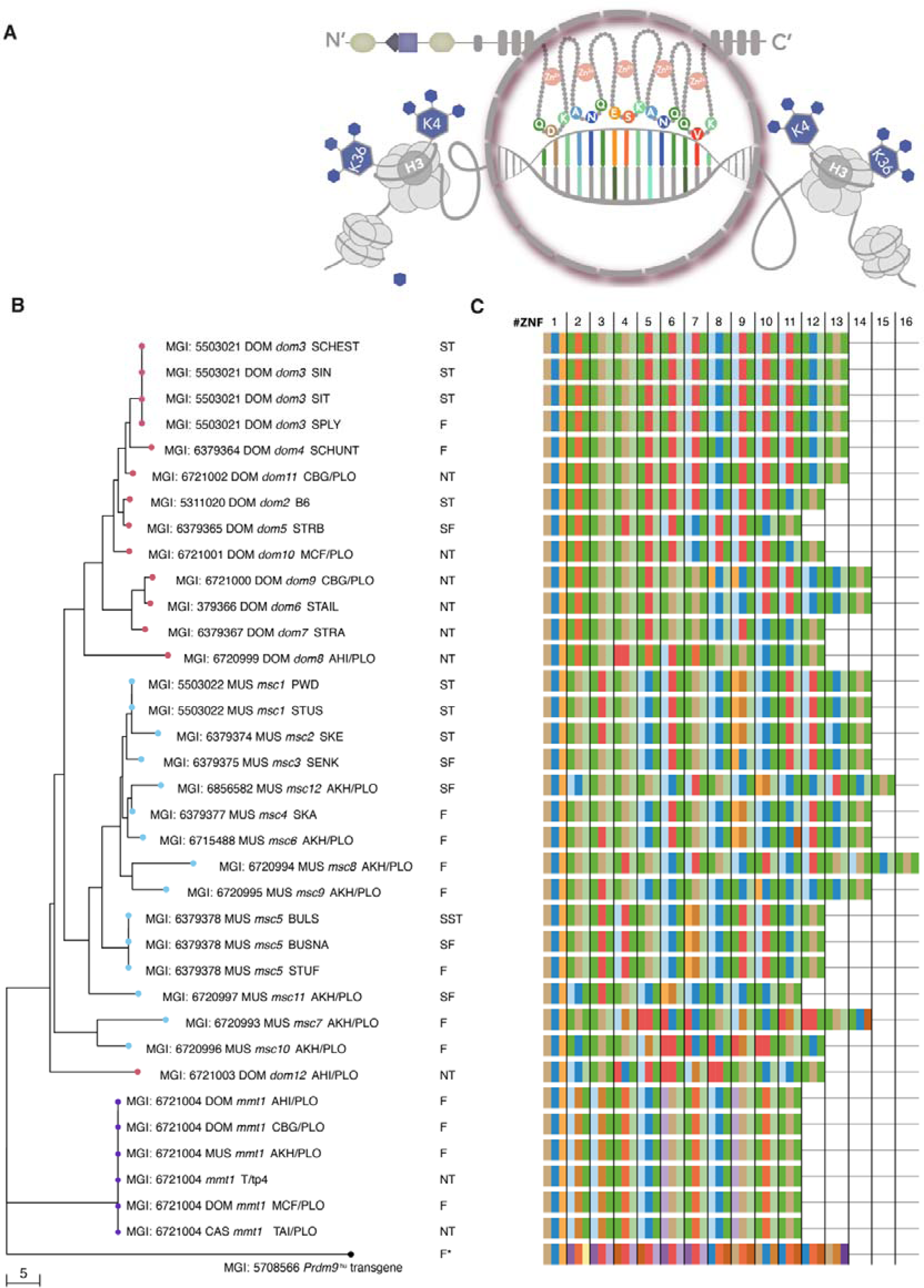
Neighbor-joining tree of the *Prdm9* exon12 minisatellite which encodes the DNA binding domain of the PRDM9 protein. **(A)** cartoon depicting the amino-acids in positions -1, 3, and 6 of the alpha-helix of each C_2_H_2_ ZNFs that are responsible for DNA binding **(B)** Neighbor-joining tree calculated on the nucleotide sequences of all PRDM9 alleles in this study and (MUKAJ *et al*. 2020), that code for the C_2_H_2_ ZNFs array, with red nodes for DOM and blue nodes for MUS alleles, and with purple nodes depicting the t-haplotype allele found in MUS and DOM **(C)** Table depicting the C_2_H_2_ ZNF array encoded by each allele, with boxes (colored as in Figure S1 and Figure S1) representing only the amino acids responsible contacting DNA of each ZNF.

The *dom12* allele, neighboring a branch of MUS alleles, displays low divergence to the last common ancestor of MUS and DOM alleles. Except for *mmt1* and *dom12*, all alleles are separated by subspecies origin (as seen in **Figure 3** and **Figure S1**). The first node separates the *dom8* allele from the Iranian population from all other DOM alleles clustering by subspecies, and a single node leads exclusively to all tested MUS alleles (**Figure 3**), which is broadly consistent with the evolutionary history of mice, with the DOM subspecies splitting first with estimated divergence time of 0.130-0.500 MYA, followed by the CAS and MUS subspecies around 0.110 - 0.320 MYA years ago (PHIFER-RIXEY *et al*. 2020). However, according to our phylogenetic reconstruction, all previously identified hybrid sterility alleles are subspecies-specific alleles of considerable divergence from a common ancestor.

However, as loci under positive selection can influence divergence times, we calculated a second distance matrix of Hamming distances after removing the hypervariable amino acids at -1, +3, and +6 of the alpha-helix, which are responsible for DNA binding (**Figure S1A**) before computing a second phylogeny. Indeed, larger divergence times are not driven by positive selection alone. The topology of the tree changes dramatically (compare **Figure 3** with **Figure S1**) for alleles that differ outside of the positively selected sites. Some repeats possess a Tryptophan (W) residue in position -5. Tryptophan’s nonpolar, aromatic, and neutral chemical properties differ from the Arginine (R) typically found in this position, which is polar and strongly basic. Secondly, there are two types of last repeats; the rarer one possesses Arginine (R) in position 13, while the more common type contains aliphatic and nonpolar Glycine (G) (**Figure S1**). The nucleotides coding for Glycine in this position appear to be the ancestral alleles, as the same amino acid is also encoded at this position in human PRDM9, included in the genetically engineered “humanized” allele B in mice, where it can rescue sterility (DAVIES *et al*. 2016)(**Figure S1**). While the *mmt1* allele remained separated, some subspecies-specific nodes disappeared. The closely related alleles to the common ancestor of *Mus musculus Prdm9* alleles are *msc11*, *msc5*, and *msc10,* which are now neighboring DOM alleles, possibly placing their origin before a clear separation into MUS and DOM subspecies. Curiously, while the full-length sequence of the *msc7* allele had previously appeared most closely related to the *msc10* allele (**Figure 3**), it is now found neighboring the *msc2* allele from SKE/JPia, within a tree of subspecies-specific alleles (**Figure S1**). While only a few alleles with substitutions outside the hypervariable sites remained separated, pointing to longer divergence times between alleles, most alleles that differed only at hypervariable sites are found on the same branch once loci under positive selection are removed. They include, on the one hand, *msc1*, *msc4*, and *msc9* and, on the other hand, *dom6* and *dom9*. The divergence between *Prdm9* alleles thus appears predominantly driven by positive selection on the hypervariable sites.

In conclusion, the complementary phylogenetic analyses support the accelerated evolution of the hypervariable DNA binding sites of PRDM9 protein and reiterate an evolutionary history in which *Mus musculus* originated in Asia and the Middle East before dispersing across Europe. The phylogenetic analyses further support a scenario in which MUS and DOM have split recently and are still speciating but do not reveal any apparent clustering of alleles co-inducing hybrid sterility by subspecies. Admittedly, no evidence was found to support the idea that the hybrid sterility susceptible alleles (*msc1, msc1, msc5, dom2, dom3,* and *dom5*) belong to the evolutionary oldest ones closest to the common ancestor. On the contrary, the *msc1, msc2*, and *dom3* alleles are the most distal, sitting on the most distant branch of the phylogenetic tree (**Figure 3** and **Figure S1**).

### Comparative analyses of DNA binding motifs

The phylogenetic approaches are based on coding-nucleotide sequences; however, PRDM9 was identified as a candidate meiotic regulator based on a sequence motif enriched in human recombination hotspots but not in primate recombination hotspots (MYERS *et al*. 2010). To enquire into the DNA-binding motif, we predicted DNA binding motifs for each conceptually translated PRDM9-ZNF using the polynomial SVM prediction method (PERSIKOV AND SINGH 2014) which is shown in (**Figure 4A**). To enquire into whether similar coding sequences of the PRDM9 ZNF-array would predict binding to the same or highly similar motifs, we used TomTom (GUPTA *et al*. 2007). Indeed, many DNA binding motifs are highly similar to each other (**Figure 4B**). These include predicted DNA binding motifs of ZNF domains encoded by *msc1, msc2*, *msc3*, *msc6, msc9,* and *msc12* in MUS, or *dom6* and *dom9*, *dom3*, *dom4* and *dom11* in DOM. Highly similar DNA binding motifs of differing ZNF domains may be able to activate the same hotspots. Cross-activation of the same hotspot by PRDM9 variants encoded by several highly similar alleles has been observed in human hotspots (BERG *et al*. 2010; BERG *et al*. 2011). Likewise, highly similar predicted DNA-binding motifs were also enriched in contemporary mouse meiotic recombination hotspots, even in other mouse strains (SMAGULOVA *et al*. 2016). To investigate putative genome-wide targets of each predicted DNA binding motif, we used PWMScan (AMBROSINI *et al*. 2018). We then quantified how many genomic targets of predicted binding sites would be shared between PRDM9 ZNF domains of different alleles. Considerable overlap in the distribution of genome-wide putative binding sites is seen particularly across highly similar alleles (**Figure 4C**). Exceptionally high overlap of putative genomic binding sites is observed for ZNF domains encoded by MUS HS alleles *msc1* and *msc2*, *msc3*, as well as between *msc4* and *msc12*, and to a lesser extent between *msc6* and *msc1*.

**Figure 4.**
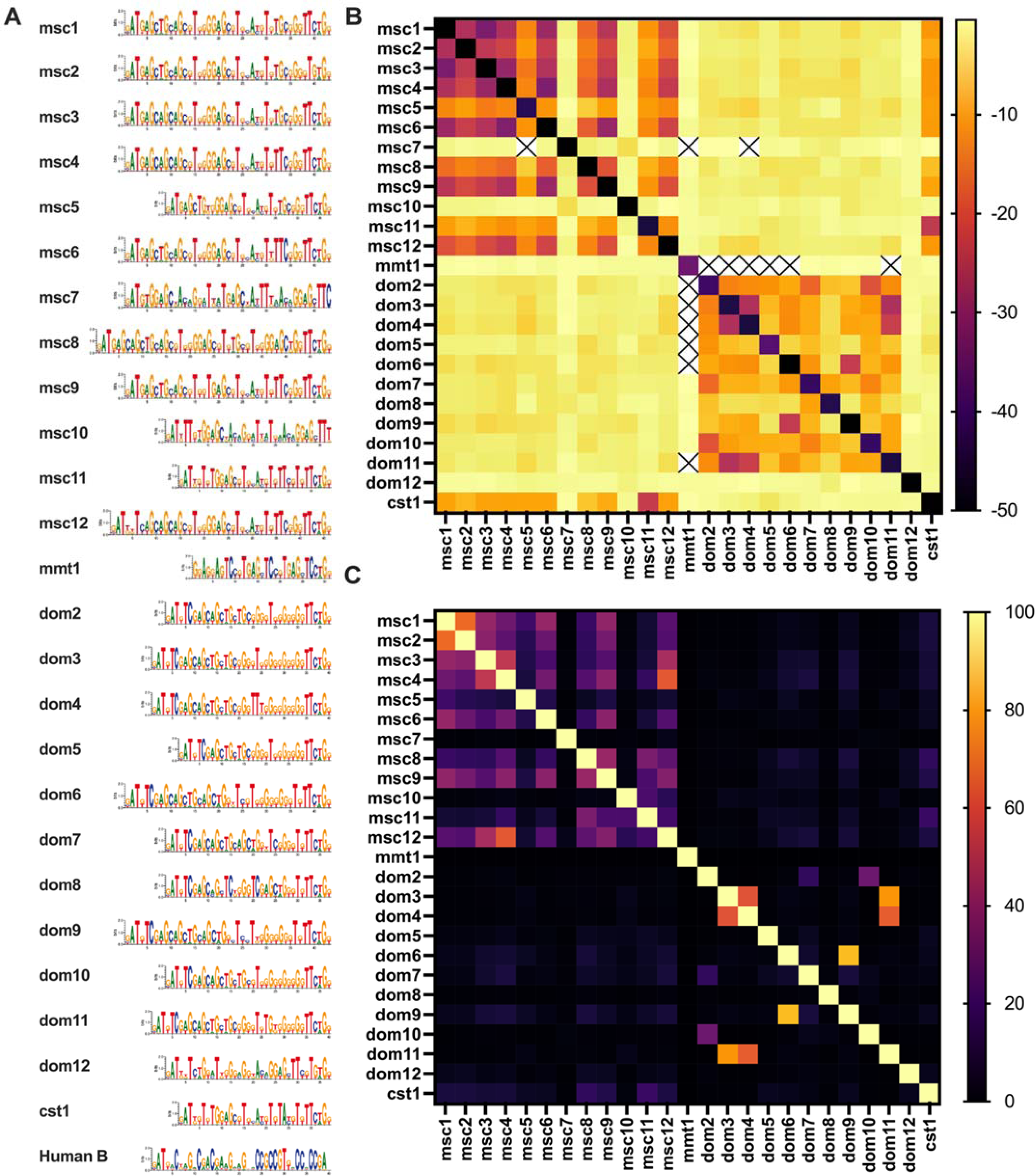
In-silico predicted PRDM9 DNA binding. **(A)** PRDM9 DNA binding motifs are represented as sequence logos of the underlying positional weight matrices, which were predicted using the Polynomial Kernel method by (PERSIKOV et al. 2009; PERSIKOV AND SINGH 2014) on translated nucleotide sequences of alleles in this study and in (MUKAJ et al. 2020) **(B)** Motifs were compared using TomTom, within the MEME suite, which computes the probabilities that a random motif would be better matched than the input motif. TomTom output P-values were log-transformed and are shown in a heat matrix, such that darker colors represent better matching of sequence motifs, and lighter colors represent weaker similarities of motifs, with crossed out values representing incidences where not similarity was found. Black boxes represent incidences when the probability of another motif binding better than the motif itself is zero, as values of zero cannot be log -transformed. **(C)** Overlap of genomic binding sites between all ZNF domain encoding alleles, which we predicted all putative binding sites for each PRDM9 binding motif genome-wide, with brighter color showing higher genomic binding site overlap between different DNA binding motifs.

## Conclusion

In summary, none of the seven novel allelic combinations of wild MUS *Prdm9* alleles with DOM *dom2* allele produced completely sterile F_1_-hybrid male offspring, consistent with the low incidence of sterile hybrids reported in the wild (TURNER *et al*. 2012). Instead, we saw either completely fertile intersubspecific hybrids (*dom2* in combination with *msc6, msc7, msc8, msc9, or msc10*) or a significant reduction of *Prdm9*-dependent fertility and increased levels of meiotic asynapsis (*dom2* in combination with *msc11,* or *msc12*). Thus, combined with the previous data from wild-derived inbred lines (MUKAJ *et al*. 2020) it appears that sterility alleles of *Prdm9* may be rare. While the data on *Prdm9* polymorphism in the wild house mouse populations are accumulating, and indeed, the *Prdm9* genes show remarkable natural allelic divergence, with more than 150 alleles having been found in mouse populations to date (BUARD *et al*. 2014; KONO *et al*. 2014; VARA *et al*. 2019; MUKAJ *et al*. 2020), little is known about their DNA binding motifs and their degree of erosion. In this context, the finding that five populations with fertile *Prdm9* alleles carried evidence of eroded *msc1* hotspots was surprising, suggesting a decoupling of the evolutionary dynamics of the PRDM9 zinc-finger domains and their binding sites. In other words, the erosion of the *msc1* hotspots may be much more common in natural populations than the *msc1* allele itself. This would align with the observation that recombination maps based on linkage disequilibrium (LD) analyses revealed a significant overlap of historical hotspots of the AKH population with contemporary *msc1-activated* hotspots of the PWD strain (WOOLDRIDGE AND DUMONT 2022). At the same time, there is weak conservation of recombination maps at the broad and the fine scale, and most hotspots are unique to the AHI, MCF, CGB, and AKH populations (WOOLDRIDGE AND DUMONT 2022). The other unanswered question relates to the evolutionary age of the *Prdm9* sterility alleles. In a simple scenario, the sterility-inducing alleles would be expected to be the oldest, situated closest to the common ancestor on the phylogenetic tree. Still, the analysis revealed the opposite: *msc1, msc2,* and *dom3* are the most distal, and therefore most likely the youngest, alleles. Another question concerns the enigmatic *t-*haplotypes present in all three major mouse subspecies and carrying, in all examined cases, the same *Prdm9* allele coding for the same zinc-finger domain. Is it so old because it arose before the ancestor split into the three subspecies? If so, why does it not behave as a sterile allele? Where did it come from if it is a recent introgression due to extremely high transmission distortion? The structure of ZNF and other sequences of *t* haplotypes shows no similarity to any extant subspecies. Clearly, more experimental evidence is needed to understand the evolutionary dynamics of PRDM9-driven hybrid sterility.

## Availability of data and materials

Nucleotide sequences of *Prdm9* alleles were deposited to Genbank under accession numbers OQ055171 - OQ055188. The fertility datasets generated and analyzed during the current study, DNA binding predictions of C_2_H_2_ zinc-finger domains encoded by each allele, as well as their genome-wide DNA binding predictions, are available in a <u>Dryad repository</u> under <u>DOI 10.5061/dryad.bzkh189cm</u>. The R package to calculate the genetic distance between complex repeats is available at https://gitlab.gwdg.de/mpievolbio-it/repeatr.

## Acknowledgments

We are grateful to Peter Donnely for the transgene *Prdm9^tm1.1^*(PRDM9)*^Wthg^* (“humanized”) strain and Attila Toth for the HORMAD2 antibody. We thank Christine Pfeifle and the entire mouse house team at the MPI in Plön for their help with mouse breeding and maintenance. We are grateful to Heike Harre for help with fertility phenotyping, Vladana Fotopulosova for help with cytology, and Nicole Thomsen for help with DNA extraction and genotyping.

## Author contributions

LOH, KFNA, ED, KKU, and AM acquired, analyzed, and interpreted data. JF, EP, and LOH conceived the project, designed the work, supervised data acquisition, and analyzed and interpreted the data. LOH wrote the manuscript. All authors have read and approved the final manuscript.

## Supplementary Data

**Figure S1:**
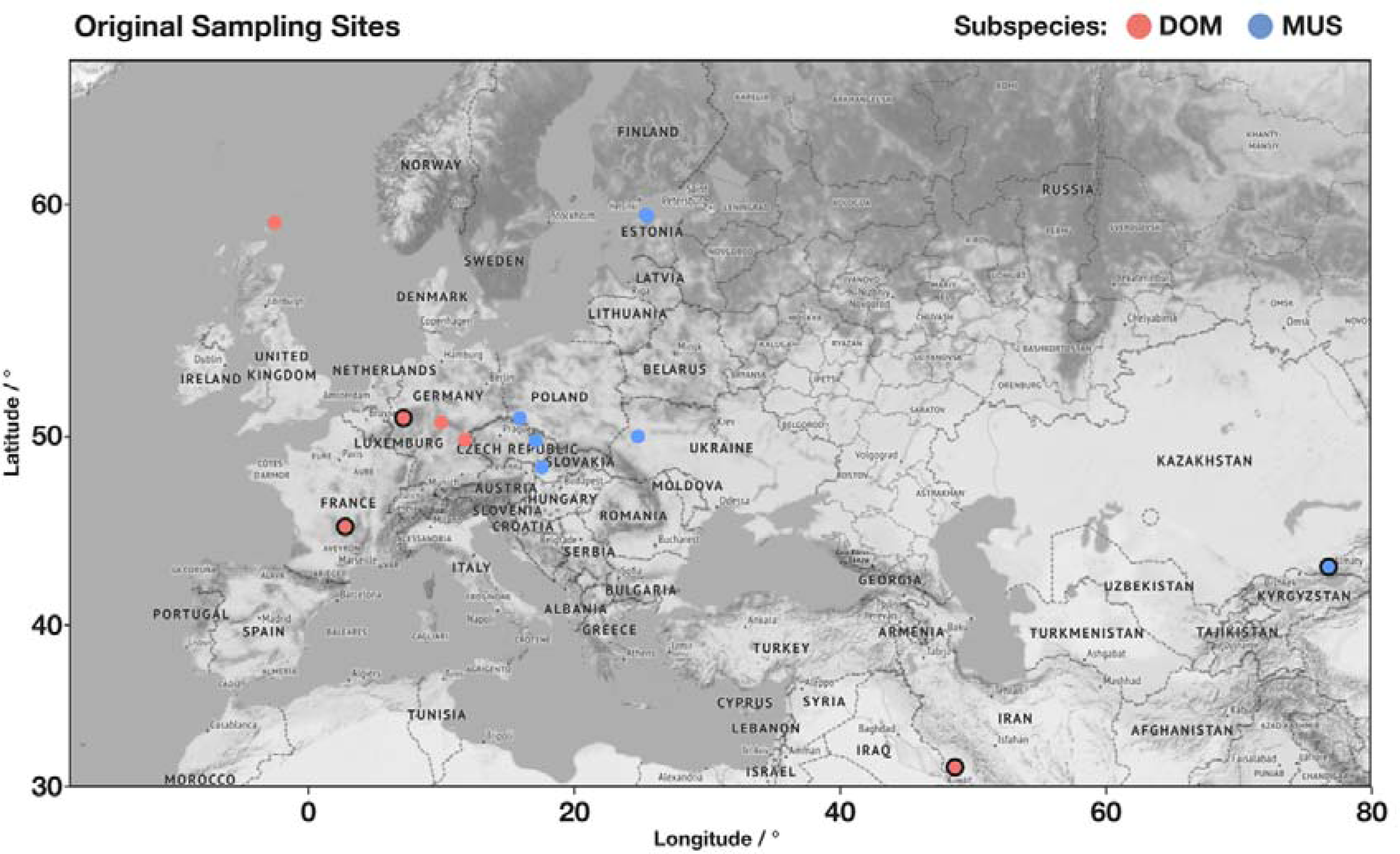
**Distribution of original sampling sites** of MUS (blue) and DOM (red), with framed circles representing mice evaluated for hybrid sterility in this study, and plain circles from (MUKAJ *et al*. 2020)

**Figure S2:**
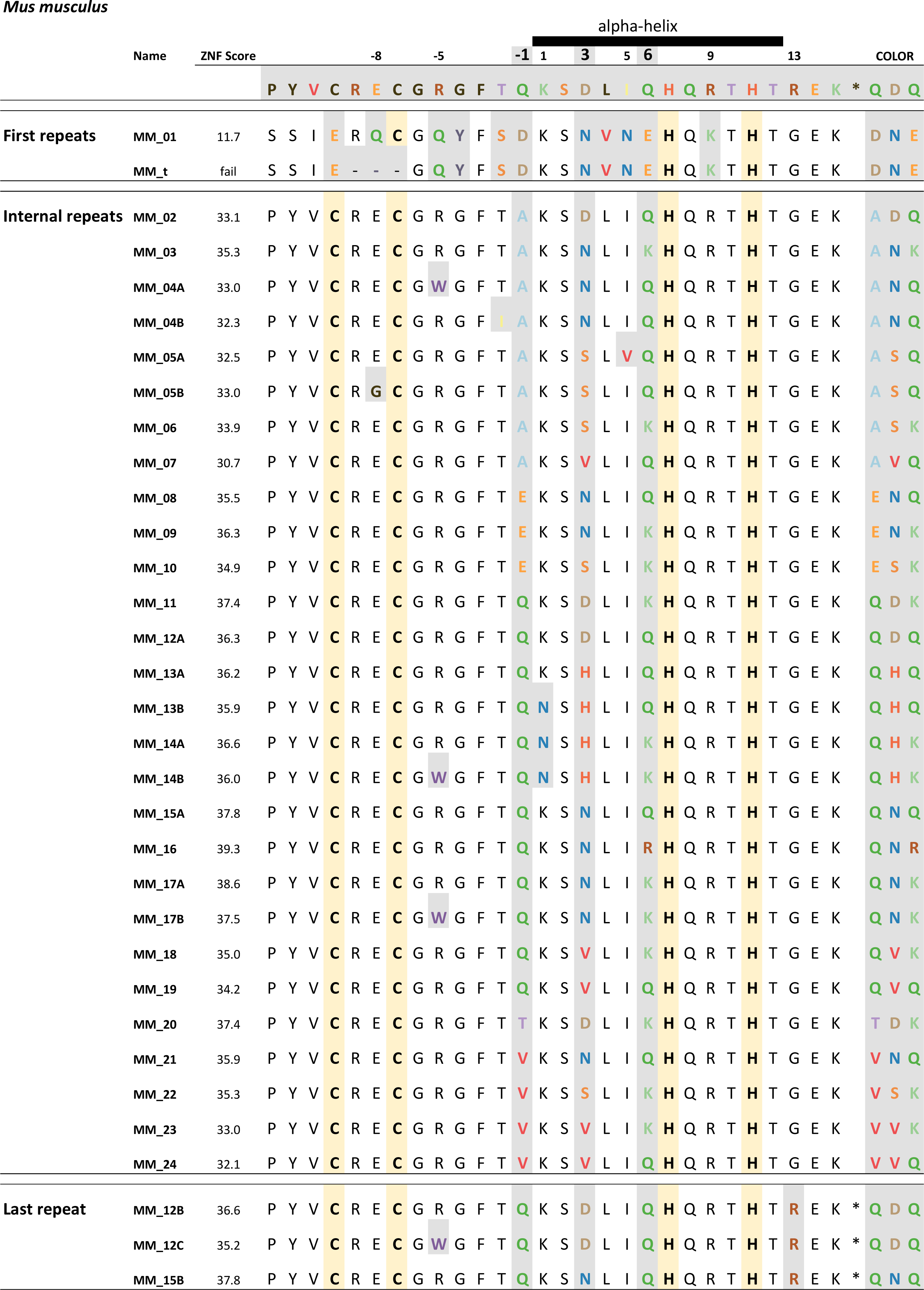
**Types of Cysteine-2-Histidine-2 (C_2_H_2_) ZNFs found in PRDM9 MUS and DOM** in this study (MUKAJ *et al*. 2020). We highlighted the Cysteines and Histidine residues in yellow and shaded all variable amino acids in gray, distinguishing between amino acids by color. As amino acids in positions -1, 3, and 6 of the alpha-helix are those responsible for the DNA-binding specificity of a given ZNF (as shown in the cartoon of **Figure 3A)**, we used them as acronyms on the right. Boxes of the same color characterize the hypervariable positions of each C_2_H_2_ zinc finger in **Figure S1** and in the phylogenetic analyses in **Figure 3**.

**Figure S3:**
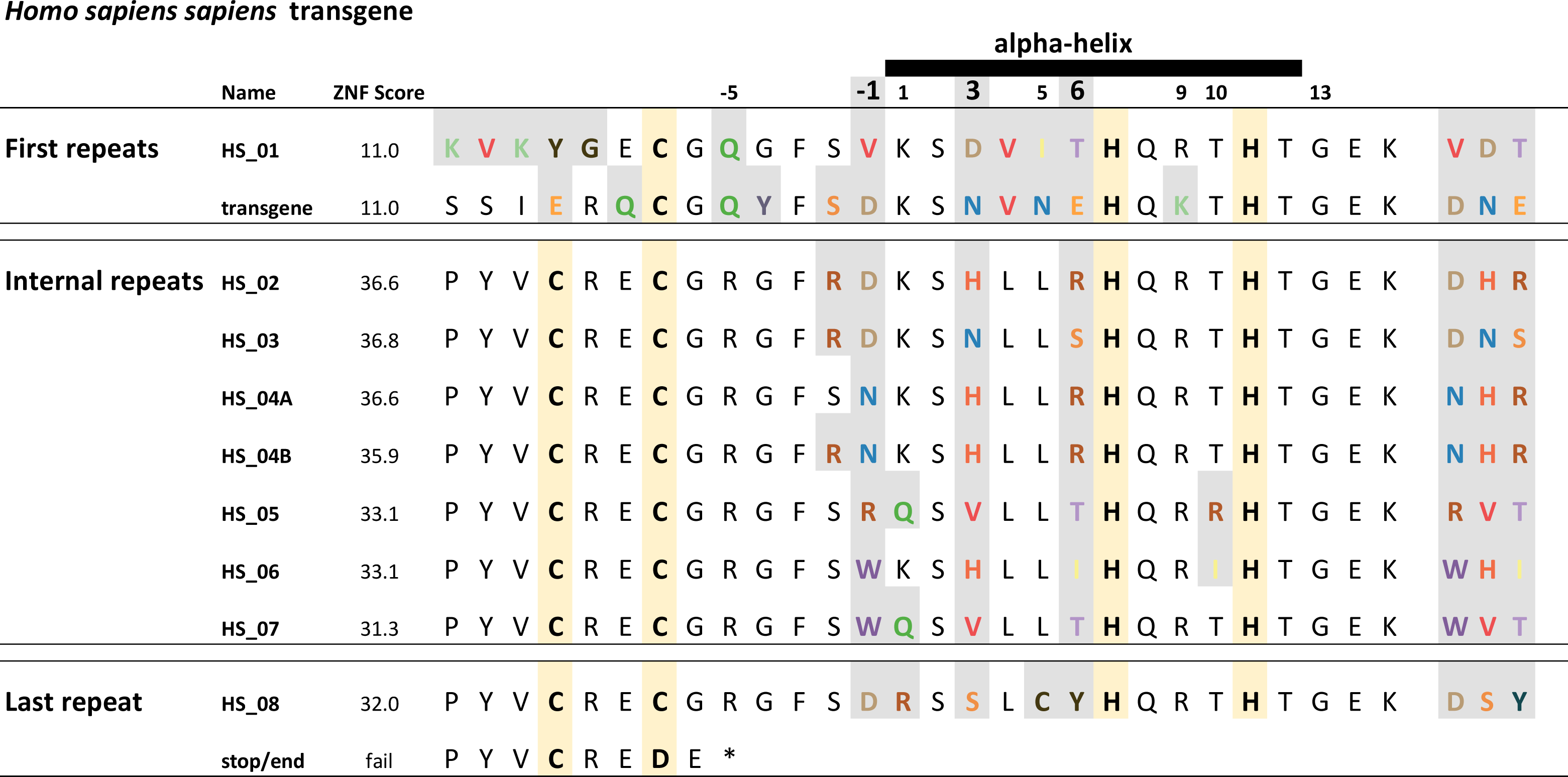
Types of C_2_H_2_ zinc fingers found in “humanized” PRDM9 variant “B.” We highlight the Cysteines and Histidine residues in yellow and shaded variable amino acids in gray. Those in positions -1, 3, and 6 of the alpha-helix are responsible for DNA binding (as shown in **Figure 3A**), and we used them as acronyms on the right. Boxes of the same color characterize the hypervariable positions of each C_2_H_2_ zinc finger in **Figure S1** and in the phylogenetic analyses in **Figure 3**.

**Figure S4:**
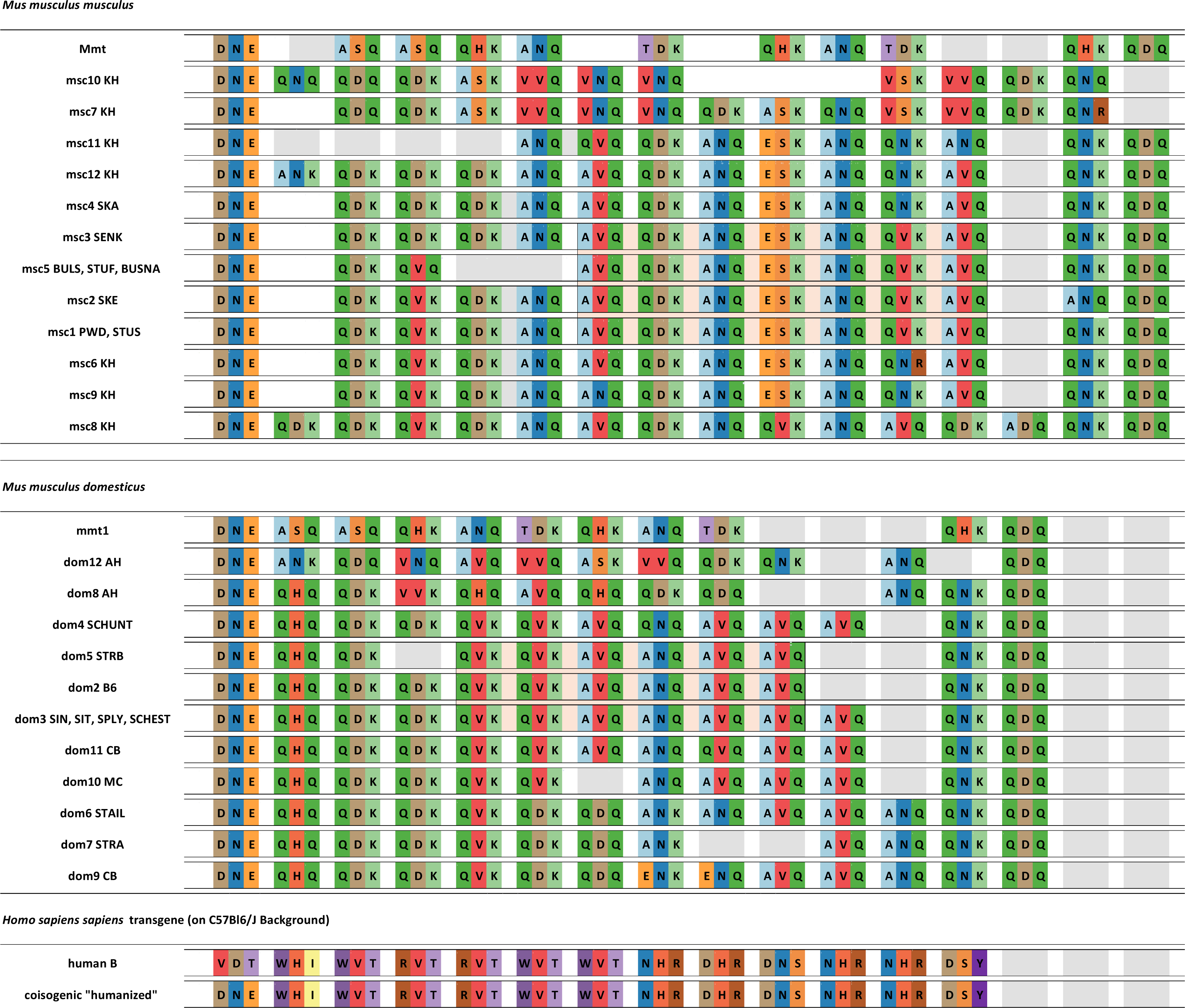
Types of C_2_H_2_ zinc finger arrays studies for hybrid sterility phenotypes. Representation of all C_2_H_2_ zinc finger arrays using only acronyms of the amino-acid positions responsible for DNA binding (as in **Figure S1** and **Figure S1**) Zinc finger arrays of *Prdm9* alleles *msc2*, *msc3*, *msc4*, msc5, *dom3, dom4, dom5, dom6 and dom7* and are from (MUKAJ *et al*. 2020). Boxes with the same colors are used to depict the hypervariable positions of each C_2_H_2_ zinc finger in the phylogenetic analyses in **Figure 3**.

**Figure S5:**
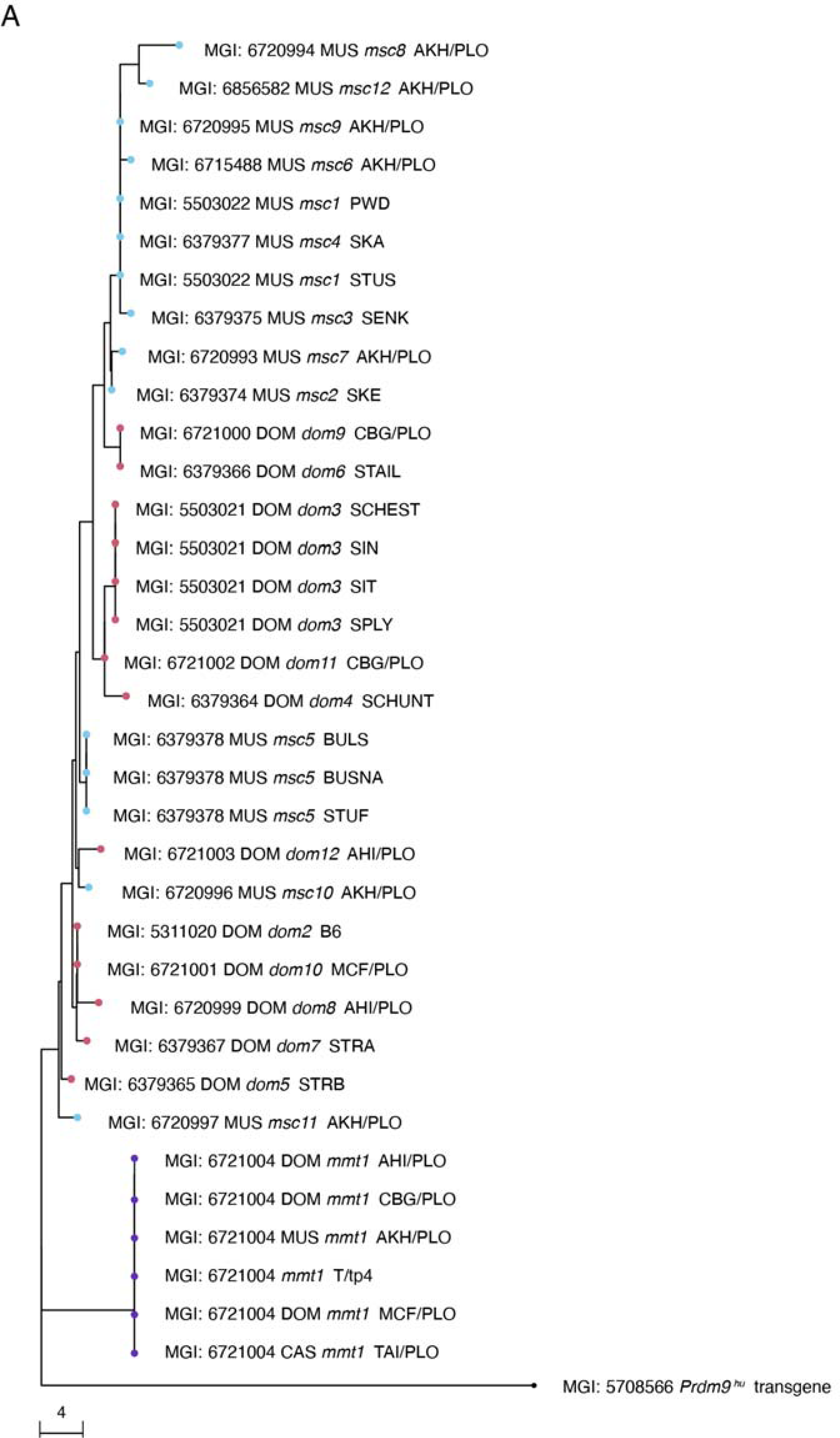
Neighbor-joining tree based on Hamming-Distances of nucleotides in the *Prdm9* minisatellite calculated after removing hypervariable nucleotides that code for the amino acids responsible for DNA binding specificity of PRDM9 and that are known to be under strong positive selection.

**Figure S6:**
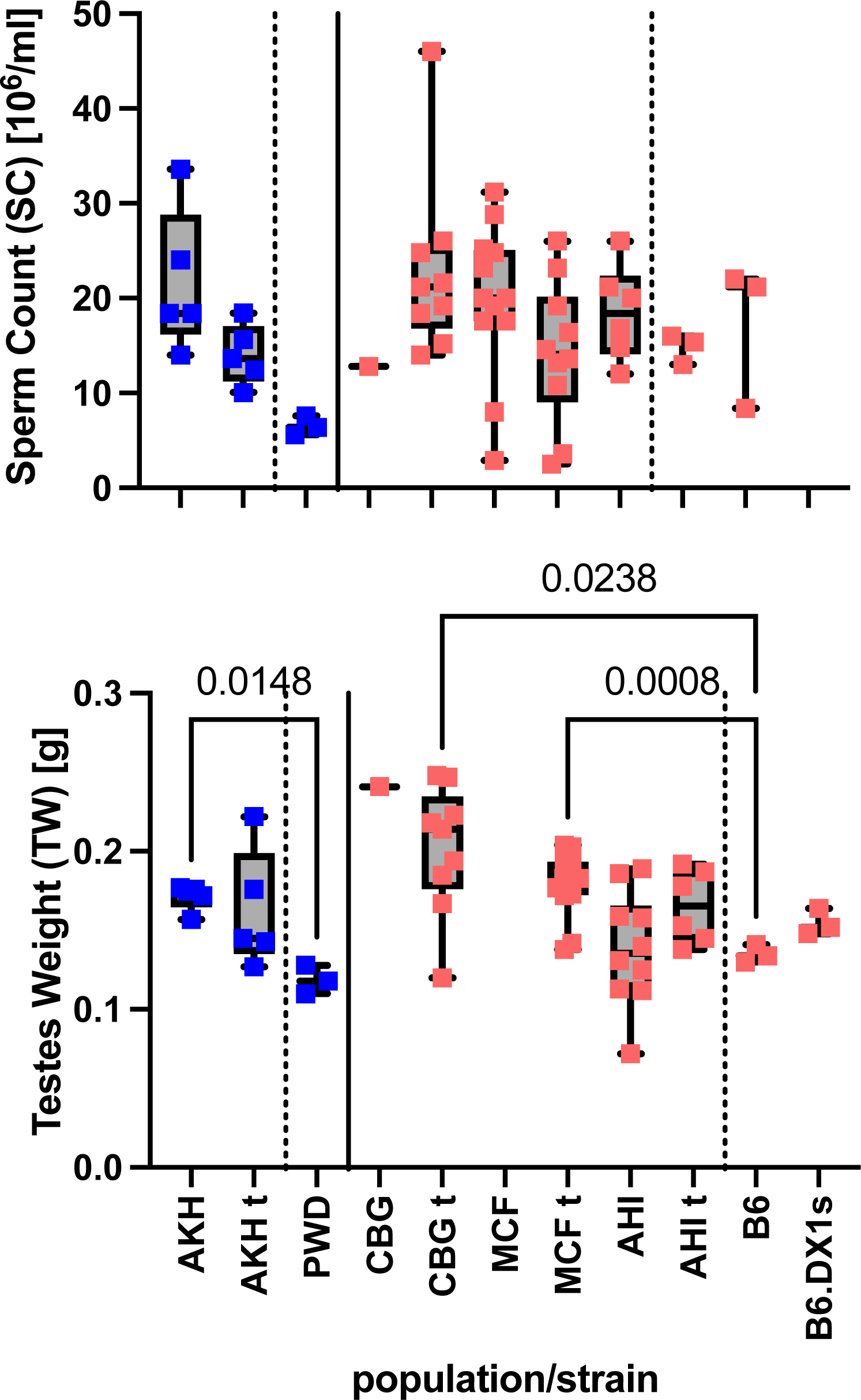
Fertility Parameters of wild mice populations in Ploen, Germany. **MUS**; **AKH** Almaty, Kazakhstan; **DOM**; **AHI** Ahvaz, Iran; **CBG** Cologne-Bonn Germany; **MCF** Massif-Central France; **t/wt** *t^t/wt^*-haplotype genotype. Statistics as described previously.

**Figure S7:**
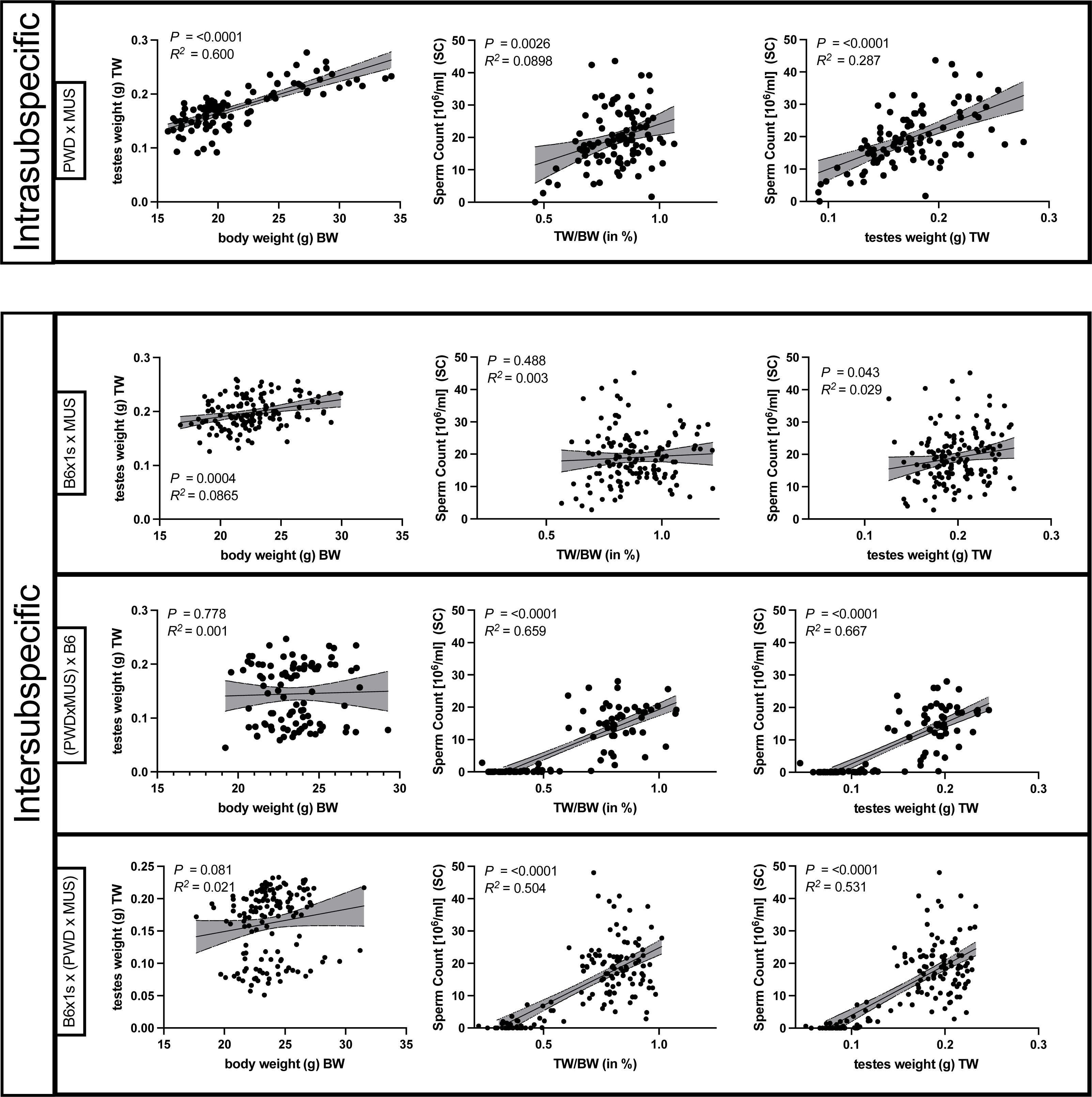
Correlation analyses of fertility parameters in all types of performed crosses. The type of crossing scheme is **(A)** Intrasubspecific cross and different **(B)** Intersubspecific, with we tested the relationship of paired testes weight to body weight, as well as the sperm count to TW/BW ratio, and testes weights (from left to right),

**Figure S8:**
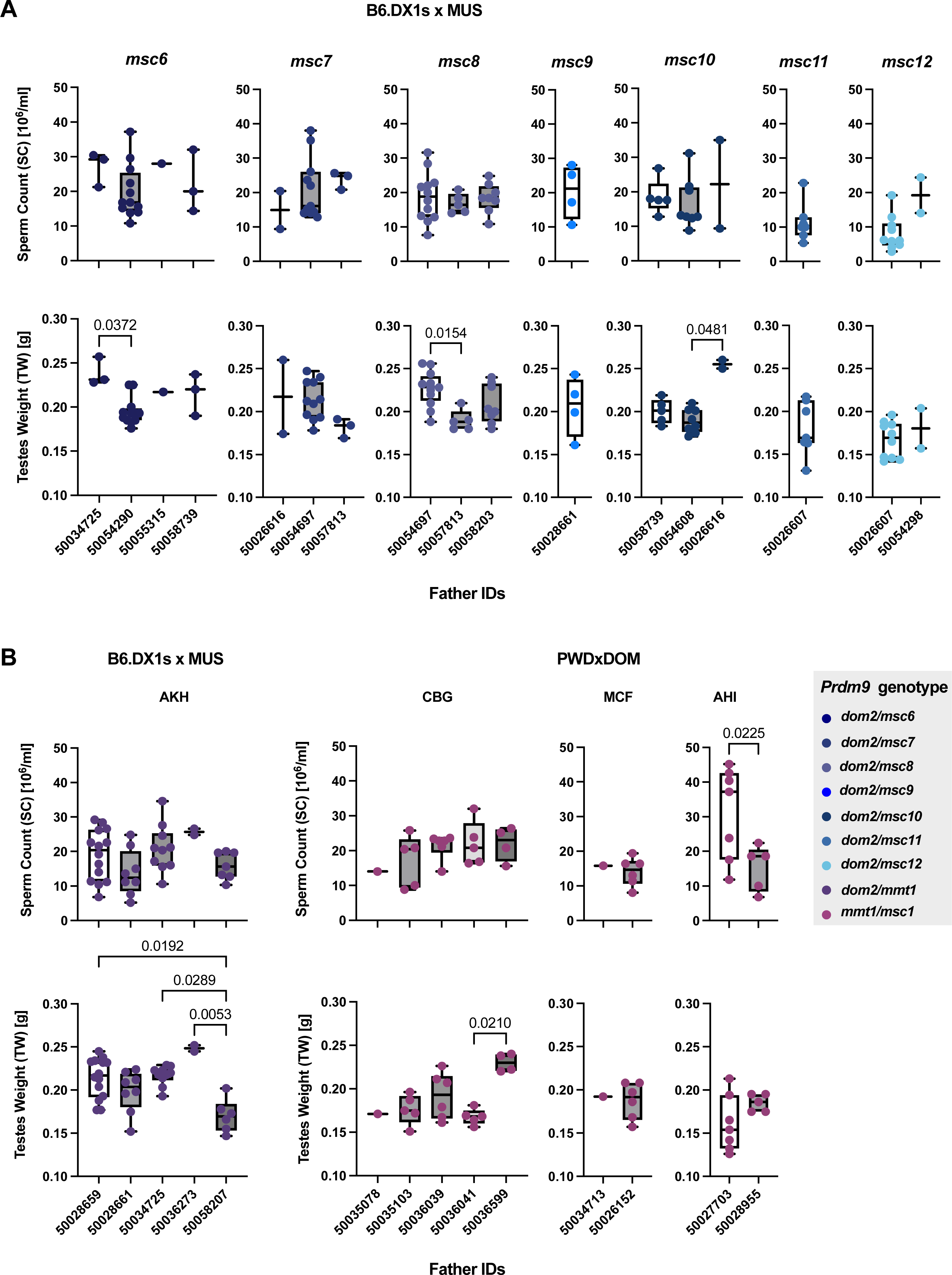
Fertility phenotypes of hybrid offspring, grouped by *Prdm9* genotype and sire ID. **(A)** MUS from Kazakhstan without *t*-haplotypes, all offspring are sorted by *Prdm9* genotype, with the father (sire) ID on the X-axis. One sire (50054290) was homozygous for *Prdm9*, all others heterozygous for different *Prdm9* alleles **(B)** Offspring with *t*-haplotypes, grouped by source population of the sires, and sire IDs on the X-axis. (left) MUS from Kazakhstan (right), DOM from Cologne-Bonn, Germany (CBG), Massif-Central, France (MCF) and Ahvaz, Iran (AHI). Data pairs were compared using Welch’s t-test, and multiple comparisons were performed using ANOVA with Kruskal-Walli’s test, corrected for multiple comparisons using Dunn’s test. Only significant values are shown on the graph with P<0.0332 (*), P<0.0021(**), P<0.0002(***), P<0.0001(****)

**Figure S9:**
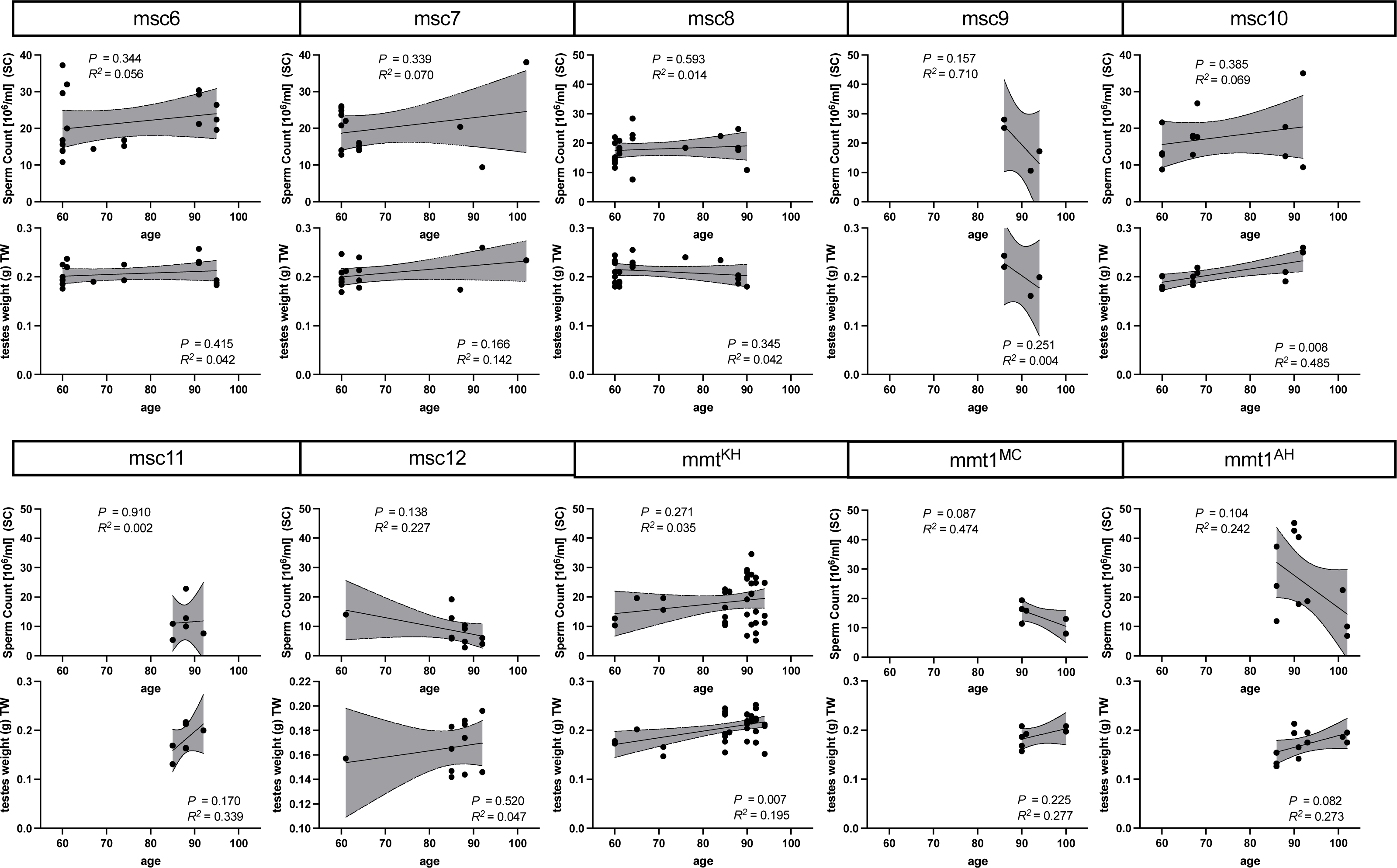
*Fertility parameters grouped by paternal Prdm9 allele from Figure 1*, analyzed by age.

**Figure S10:**
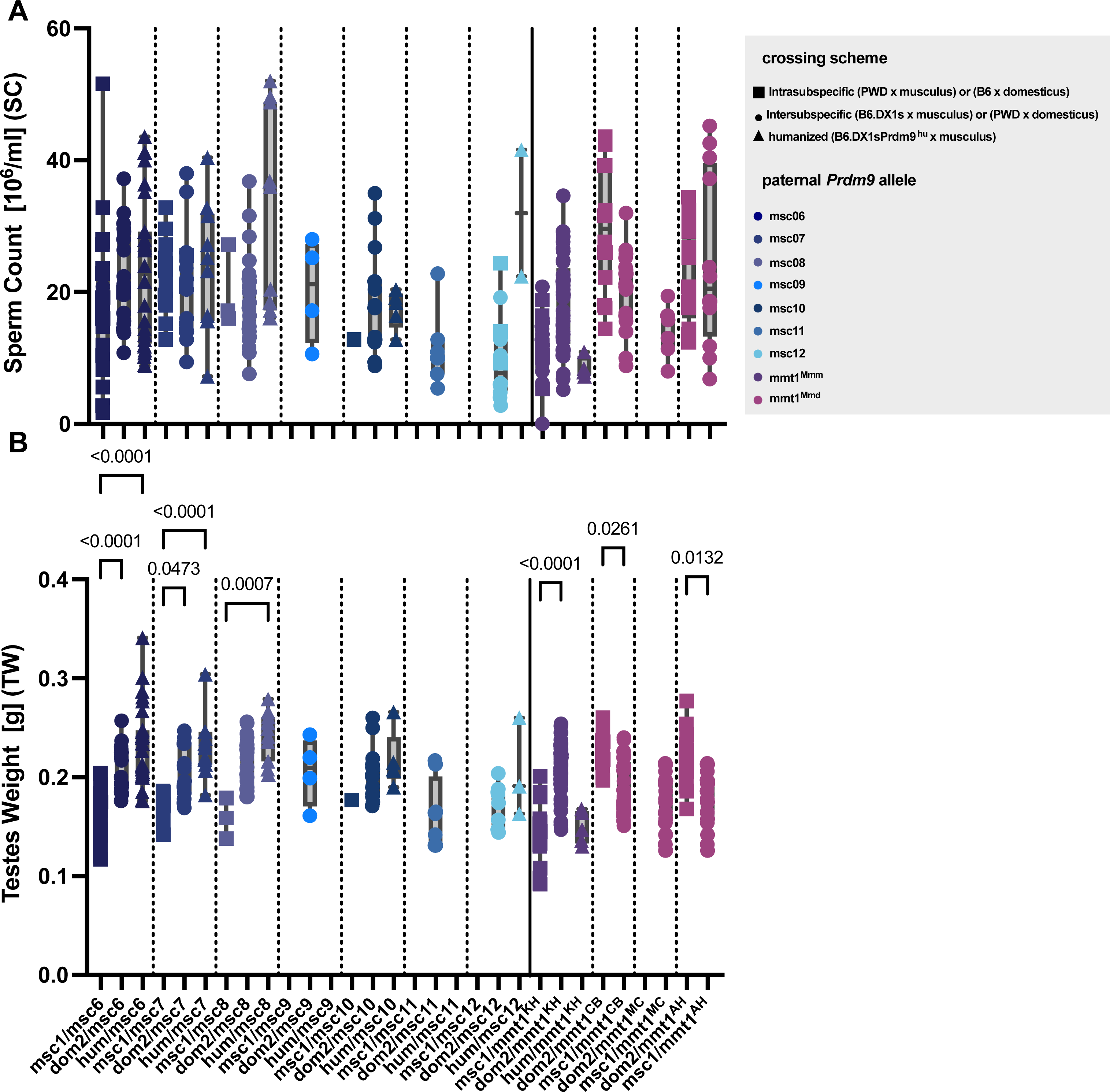
Fertility parameters of intra- and intersubspecific F_1_-hybrids. **(A)** sperm count **(B)** paired testes weight of intersubspecific and Intrasubspecific mice. Paternal *Prdm9* alleles are distinguished by colors, while the shape of the data points distinguishes crossing schemes. (Square data points) Intrasubspecific male F_1_ offspring of PWD females crossed to MUS males, or Intrasubspecific B6 females crossed to DOM males. Two types of interspecific crosses were performed; MUS males were crossed to B6.DX1s females and wild DOM males were crossed to PWD females (circles as data points, also shown in **Figure 1**); for the MUS males, a third cross was performed where they were mated with B6.DX1s .*Prdm9^hu^*females (triangular data points). Since most males used as sires were heterozygous for different allelic combinations of *Prdm9,* data from several crosses pooled by *Prdm9* genotype of the F_1_-hybrid offspring, and the significance of differences in fertility phenotypes was evaluated using ordinary one-way ANOVA with Bonferroni correction as before.

**Figure S11:**
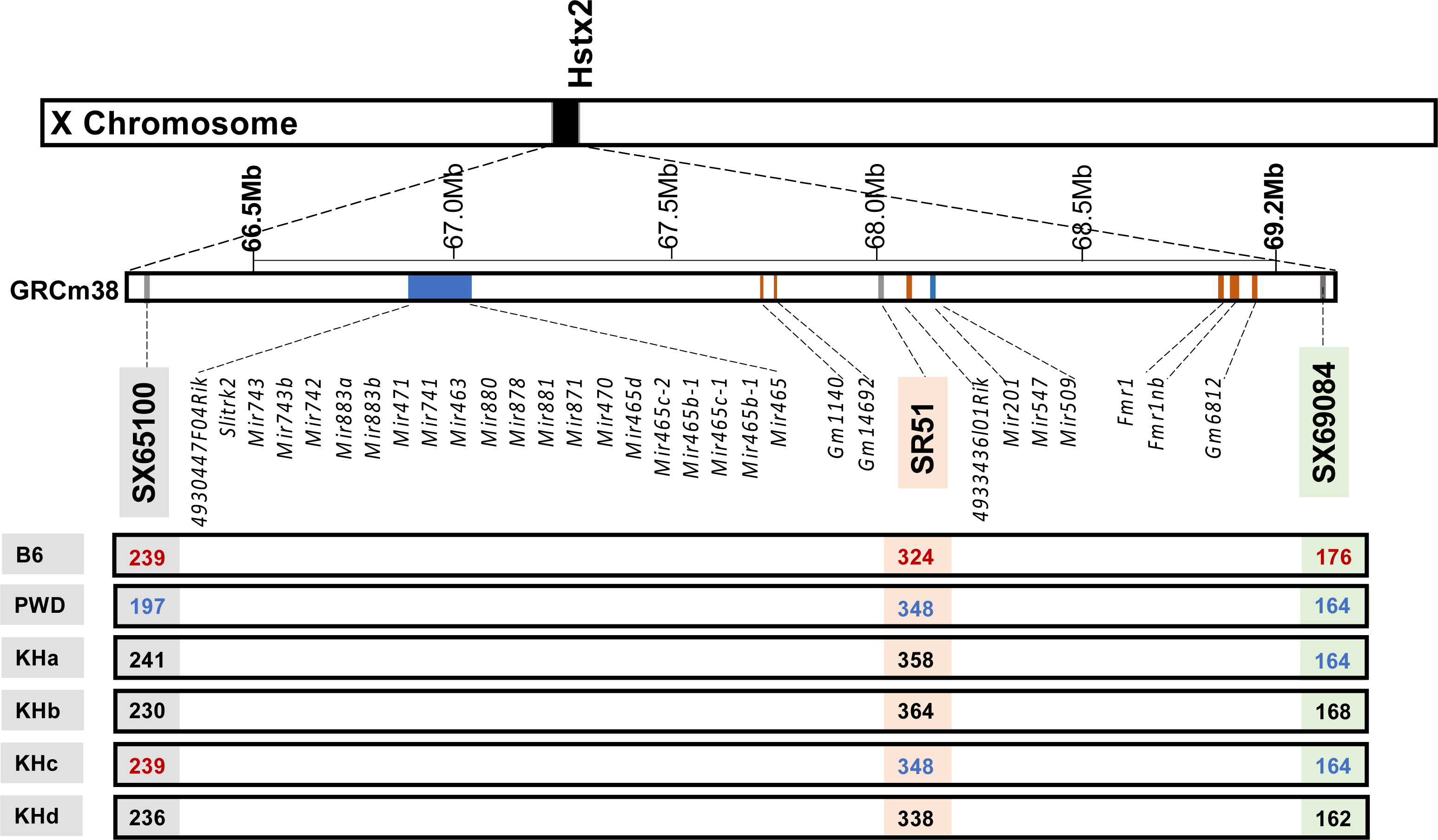
**X-chromosomal haplotypes across the refined *Hstx2* locus** from (Lustyk et al. 2019) based on allelic variation at microsatellite markers SX65100, SR51, and SX69084 in the laboratory and wild mice. All genes and microRNAs annotated within this locus are shown in the mm38 reference genome.

**Figure S12:**
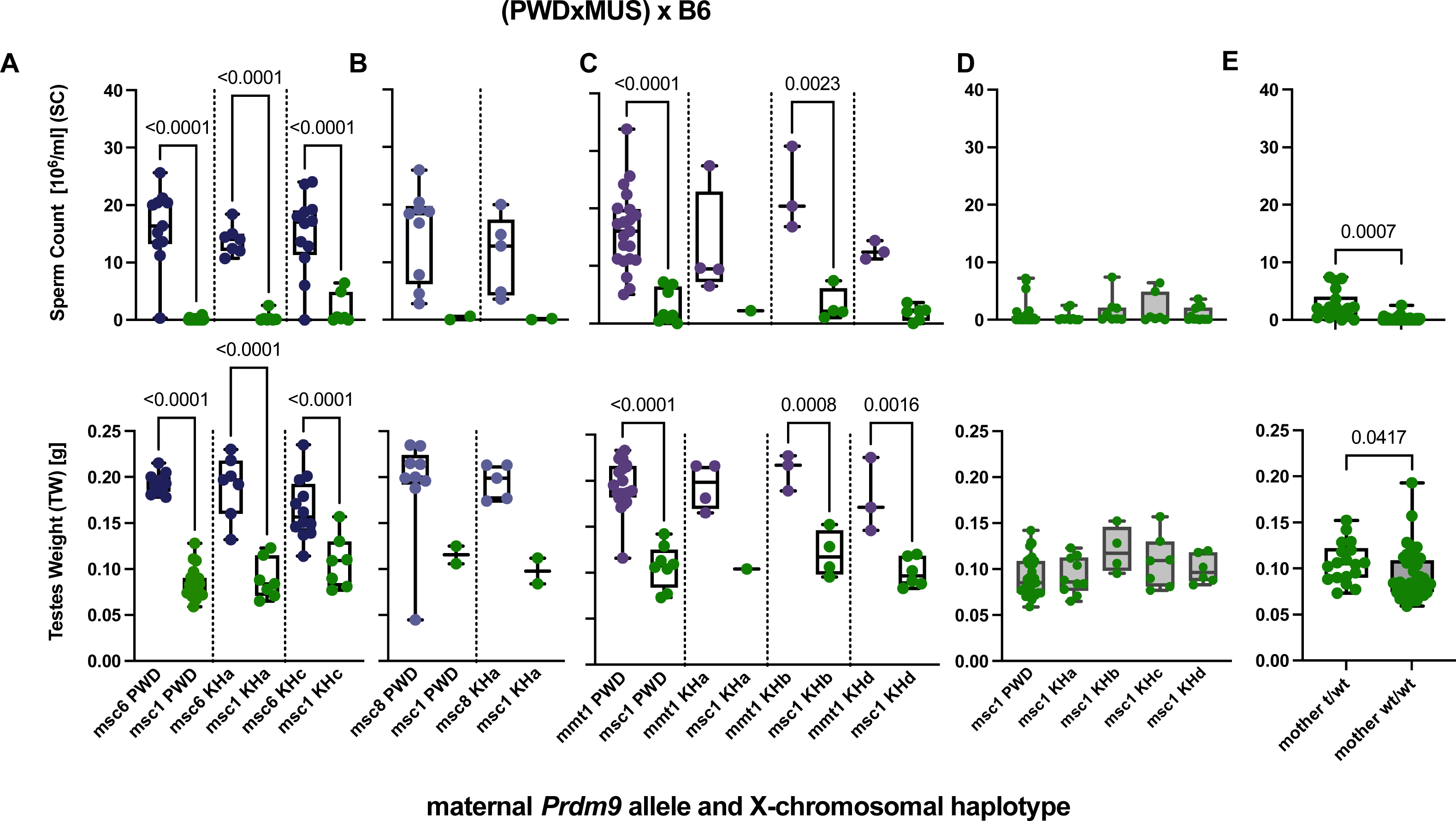
Segregation analyses of maternal *Prdm9* genotype together with the variation of wild X-chromosomal haplotypes. Intersubspecific F_1_ male offspring of Intraspecific MUS hybrid females crossed to B6 males (as in **Figure 2B**) that inherited wild MUS X-chromosomal haplotypes. X-axes depict both the maternal *Prdm9* allele and wild MUS X-chromosomal haplotype, defined as shown in (**Figure S1**). Fertility parameters, sperm count (top), and paired testes weights (bottom) are shown, and the significance of differences in fertility phenotypes was evaluated as before. (A-C) Segregation analyses grouped by maternal *Prdm9* allele combination (A) maternal *msc1* and *msc6* alleles (B) maternal *msc1* and *msc8* alleles (C) *msc1* and *mmt1^KH^* alleles (D-E) Pooled data of hybrids from different mothers that all inherited the same *msc1/dom2* allelic combination (D) pooled by X-chromosomal haplotype (E) pooled by whether mothers possessed t-haplotypes.

**Table S1:** Variation, Introgression, and Recombination rate of outcrossed populations with data from (a) (LAWAL *et al*. 2021) (b) (WOOLDRIDGE AND DUMONT 2022) (c) (BANKER *et al*. 2022) (d) (STAUBACH *et al*. 2012)

| outcrossed populations | # SNPS | SNP density (bp/SNP) | # population private variants | Average introgression per individual (%) | % of the genome affected by introgression |
| --- | --- | --- | --- | --- | --- |
| AKH | 10,937,288<sup>a</sup> | 225<sup>b</sup> | 1,872,782<sup>a</sup> | NA | 4.90<sup>d</sup> |
| MCF | 11,108,085<sup>a</sup> | 222<sup>b</sup> | 2,208,483<sup>a</sup> | 0.069<sup>c</sup> | 2.80<sup>d</sup> |
| CBG | 11,930,888<sup>a</sup> | 206<sup>b</sup> | 545,881<sup>a</sup> | 0.068<sup>c</sup> | 2.80<sup>d</sup> |
| AHI | 17,877,283<sup>a</sup> | 138<sup>b</sup> | 3,333,440<sup>a</sup> | 0.036<sup>c</sup> | NA |

**Table S2.**
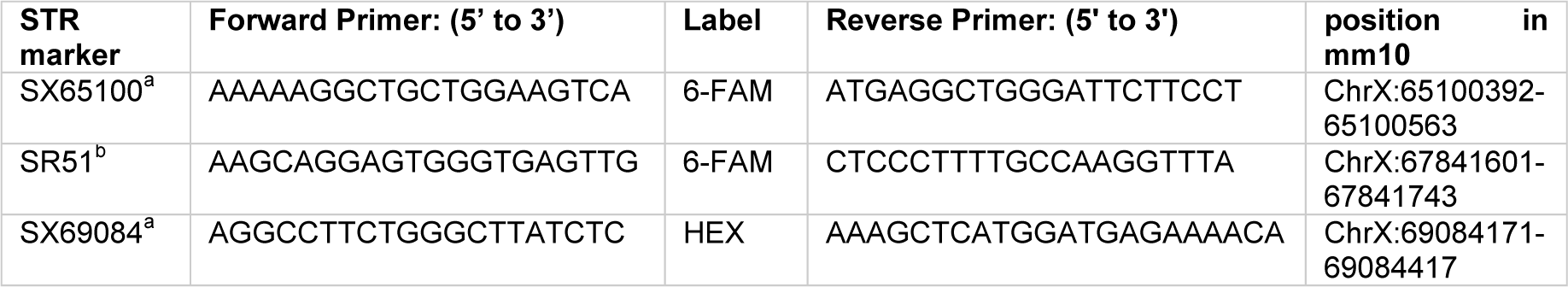
Primers used to genotype the *Hstx2* interval from (a)(Lustyk et al. 2019) and (b) P. Jansa personal communication.

**Table S 3.** Difference in Sperm count (SC) grouped by *Prdm9* Genotype of F_1_ hybrid male offspring, evaluated using Ordinary one-way ANOVA with Bonferroni’s multiple comparisons test.

| Ordinary one-way ANOVA | Summary | Bonferroni Adjusted -P-Value | Mean 1 | Mean 2 | Mean Diff. |
| --- | --- | --- | --- | --- | --- |
| *msc6* vs. *msc1* | *** | **0.0001** | 22.1 | 0 | 22.1 |
| *msc7* vs. *msc1* | *** | **0.0007** | 20.8 | 0 | 20.8 |
| *msc8* vs. *msc1* | ** | **0.0043** | 18.2 | 0 | 18.2 |
| *msc9* vs. *msc1* | * | **0.0246** | 20.3 | 0 | 20.3 |
| *msc10* vs. *msc1* | ** | **0.0088** | 18 | 0 | 18 |
| *msc11* vs. *msc1* | ns | >0.9999 | 11.4 | 0 | 11.4 |
| *msc12* vs. *msc1* | ns | >0.9999 | 9.97 | 0 | 9.97 |
| *mmt1<sup>KH</sup>* vs. *msc1* | ** | **0.0029** | 18.2 | 0 | 18.2 |
| *msc6* vs. *msc12* | *** | **0.0005** | 22.1 | 9.97 | 12.1 |
| *msc7* vs. *msc12* | ** | **0.0099** | 20.8 | 9.97 | 10.8 |
| *msc8* vs. *msc12* | ns | 0.1046 | 18.2 | 9.97 | 8.26 |
| *msc9* vs. *msc12* | ns | >0.9999 | 20.3 | 9.97 | 10.3 |
| *msc10* vs. *msc12* | ns | 0.3438 | 18 | 9.97 | 8.06 |
| *msc11* vs. *msc12* | ns | >0.9999 | 11.4 | 9.97 | 1.39 |
| *mmt1<sup>KH</sup>* vs. *msc12* | * | **0.0464** | 18.2 | 9.97 | 8.28 |
| *mmt1<sup>CB</sup>* vs. *dom2* | **** | **<0.0001** | 20.3 | 0 | 20.3 |
| *mmt1<sup>MC</sup>* vs. *dom2* | ns | 0.1394 | 14.3 | 0 | 14.3 |
| *mmt1<sup>AH</sup>* vs. *dom2* | **** | **<0.0001** | 24.6 | 0 | 24.6 |

**Table S 4.** Difference in testis weight (TW) grouped by *Prdm9* Genotype of F_1_ hybrid male offspring, evaluated using Ordinary one-way ANOVA with Bonferroni’s multiple comparisons test.

| Ordinary one-way ANOVA | Summary | Bonferroni Adjusted -P- Value | Mean 1 | Mean 2 | Mean Diff. |
| --- | --- | --- | --- | --- | --- |
| *msc6* vs. *msc1* | **** | <0.0001 | 0.206 | 0.0623 | 0.144 |
| *msc7* vs. *msc1* | **** | <0.0001 | 0.208 | 0.0623 | 0.145 |
| *msc8* vs. *msc1* | **** | <0.0001 | 0.213 | 0.0623 | 0.151 |
| *msc9* vs. *msc1* | **** | <0.0001 | 0.206 | 0.0623 | 0.143 |
| *msc10* vs. *msc1* | **** | <0.0001 | 0.202 | 0.0623 | 0.139 |
| *msc11* vs. *msc1* | **** | <0.0001 | 0.18 | 0.0623 | 0.117 |
| *msc12* vs. *msc1* | **** | <0.0001 | 0.169 | 0.0623 | 0.107 |
| *mmt1<sup>KH</sup>* vs. *msc1* | **** | <0.0001 | 0.203 | 0.0623 | 0.141 |
| *msc6* vs. *msc12* | * | 0.0115 | 0.206 | 0.169 | 0.0371 |
| *msc7* vs. *msc12* | * | 0.0107 | 0.208 | 0.169 | 0.0386 |
| *msc8* vs. *msc12* | *** | 0.0002 | 0.213 | 0.169 | 0.0442 |
| *msc9* vs. *msc12* | ns | >0.9999 | 0.206 | 0.169 | 0.0365 |
| *msc10* vs. *msc12* | ns | 0.1165 | 0.202 | 0.169 | 0.0324 |
| *msc11* vs. *msc12* | ns | >0.9999 | 0.18 | 0.169 | 0.0103 |
| *mmt1<sup>CB</sup>* vs. *dom2* | **** | <0.0001 | 0.189 | 0.0705 | 0.118 |
| *mmt1<sup>MC</sup>* vs. *dom2* | **** | <0.0001 | 0.188 | 0.0705 | 0.118 |
| *mmt1<sup>AH</sup>* vs. *dom2* | **** | <0.0001 | 0.171 | 0.0705 | 0.101 |

## References

Abe, K., M. Yuzuriha, M. Sugimoto, M. S. Ko, M. Brathwaite et al., 2004 Gene content of the 750-kb critical region for mouse embryonic ectoderm lethal tcl-w5. Mamm Genome 15: 265–276.

Altemose, N., N. Noor, E. Bitoun, A. Tumian, M. Imbeault et al., 2017a Human PRDM9 can bind and activate promoters, and other zinc-finger proteins associate with reduced recombination in cis.

Altemose, N., N. Noor, E. Bitoun, A. Tumian, M. Imbeault et al., 2017b A map of human PRDM9 binding provides evidence for novel behaviors of PRDM9 and other zinc-finger proteins in meiosis. Elife 6.

Ambrosini, G., R. Groux and P. Bucher, 2018 PWMScan: a fast tool for scanning entire genomes with a position-specific weight matrix. Bioinformatics 34: 2483–2484.

Anderson, L. K., A. Reeves, L. M. Webb and T. Ashley, 1999 Distribution of Crossing Over on Mouse Synaptonemal Complexes Using Immunofluorescent Localization of MLH1 Protein. Genetics 151: 1569–1579.

Baker, C. L., S. Kajita, M. Walker, R. L. Saxl, N. Raghupathy et al., 2015 PRDM9 drives evolutionary erosion of hotspots in Mus musculus through haplotype-specific initiation of meiotic recombination. PLoS Genet 11: e1004916.

Baker, C. L., M. Walker, S. Kajita, P. M. Petkov and K. Paigen, 2014 PRDM9 binding organizes hotspot nucleosomes and limits Holliday junction migration. Genome Res 24: 724–732.

Balcova, M., B. Faltusova, V. Gergelits, T. Bhattacharyya, O. Mihola et al., 2016 Hybrid Sterility Locus on Chromosome X Controls Meiotic Recombination Rate in Mouse. PLoS Genet 12: e1005906.

Banker, S. E., F. Bonhomme and M. W. Nachman, 2022 Bidirectional Introgression between Mus musculus domesticus and Mus spretus. Genome Biol Evol 14.

Bateson, W., 1909 Heredity and Variation in Modern Lights.

Baudat, F., J. Buard, C. Grey, A. Fledel-Alon, C. Ober et al., 2010 PRDM9 is a major determinant of meiotic recombination hotspots in humans and mice. Science 327: 836–840.

Baudat, F., Y. Imai and B. de Massy, 2013 Meiotic recombination in mammals: localization and regulation. Nat Rev Genet 14: 794–806.

Berg, I. L., R. Neumann, K. W. Lam, S. Sarbajna, L. Odenthal-Hesse et al., 2010 PRDM9 variation strongly influences recombination hotspot activity and meiotic instability in humans. Nat Genet 42: 859–863.

Berg, I. L., R. Neumann, S. Sarbajna, L. Odenthal-Hesse, N. J. Butler et al., 2011 Variants of the protein PRDM9 differentially regulate a set of human meiotic recombination hotspots highly active in African populations. Proc Natl Acad Sci U S A 108: 12378–12383.

Bhattacharyya, T., S. Gregorova, O. Mihola, M. Anger, J. Sebestova et al., 2013 Mechanistic basis of infertility of mouse intersubspecific hybrids. Proc Natl Acad Sci U S A 110: E468–477.

Bhattacharyya, T., R. Reifova, S. Gregorova, P. Simecek, V. Gergelits et al., 2014 X chromosome control of meiotic chromosome synapsis in mouse inter-subspecific hybrids. PLoS Genet 10: e1004088.

Billings, T., E. D. Parvanov, C. L. Baker, M. Walker, K. Paigen et al., 2013 DNA binding specificities of the long zinc-finger recombination protein PRDM9. Genome Biol 14: R35.

Boulton, A., R. S. Myers and R. J. Redfield, 1997 The hotspot conversion paradox and the evolution of meiotic recombination. Proc Natl Acad Sci U S A 94: 8058–8063.

Brick, K., S. Thibault-Sennett, F. Smagulova, K. G. Lam, Y. Pu et al., 2018 Extensive sex differences at the initiation of genetic recombination. Nature 561: 338–342.

Buard, J., E. Rivals, D. Dunoyer de Segonzac, C. Garres, P. Caminade et al., 2014 Diversity of Prdm9 zinc finger array in wild mice unravels new facets of the evolutionary turnover of this coding minisatellite. PLoS One 9: e85021.

Cole, F., F. Baudat, C. Grey, S. Keeney, B. de Massy et al., 2014 Mouse tetrad analysis provides insights into recombination mechanisms and hotspot evolutionary dynamics. Nat Genet 46: 1072–1080.

Damm, E., and L. Odenthal-Hesse, 2022 Orchestrating recombination initiation in mice and men. Current Topics in Developmental Biology 151.

Damm, E., K. K. Ullrich, W. B. Amos and L. Odenthal-Hesse, 2022 Evolution of the recombination regulator PRDM9 in minke whales. BMC Genomics 23: 212.

Davies, B., E. Hatton, N. Altemose, J. G. Hussin, F. Pratto et al., 2016 Re-engineering the zinc fingers of PRDM9 reverses hybrid sterility in mice. Nature 530: 171–176.

de la Fuente, R., M. T. Parra, A. Viera, A. Calvente, R. Gomez et al., 2007 Meiotic pairing and segregation of achiasmate sex chromosomes in eutherian mammals: the role of SYCP3 protein. PLoS Genet 3: e198.

Dobzhansky, T., 1936 Studies on Hybrid Sterility. II. Localization of Sterility Factors in Drosophila Pseudoobscura Hybrids. Genetics 21: 113–135.

Dzur-Gejdosova, M., P. Simecek, S. Gregorova, T. Bhattacharyya and J. Forejt, 2012 Dissecting the genetic architecture of F1 hybrid sterility in house mice. Evolution 66: 3321–3335.

Eram, M. S., S. P. Bustos, E. Lima-Fernandes, A. Siarheyeva, G. Senisterra et al., 2014 Trimethylation of histone H3 lysine 36 by human methyltransferase PRDM9 protein. J Biol Chem 289: 12177–12188.

Fernandez-Capetillo, O., S. K. Mahadevaiah, A. Celeste, P. J. Romanienko, R. D. Camerini-Otero et al., 2003 H2AX Is Required for Chromatin Remodeling and Inactivation of Sex Chromosomes in Male Mouse Meiosis. Developmental Cell 4: 497–508.

Flachs, P., O. Mihola, P. Simecek, S. Gregorova, J. C. Schimenti et al., 2012 Interallelic and intergenic incompatibilities of the Prdm9 (Hst1) gene in mouse hybrid sterility. PLoS Genet 8: e1003044.

Forejt, J., 2016 Genetics: Asymmetric breaks in DNA cause sterility. Nature 530: 167–168.

Forejt, J., S. Gregorova and P. Jansa, 1988 Three new t-haplotypes of Mus musculus reveal structural similarities to t-haplotypes of Mus domesticus. Genet Res 51: 111–119.

Forejt, J., and P. Ivanyi, 1974 Genetic studies on male sterility of hybrids between laboratory and wild mice (Mus musculus L.). Genet Res 24: 189–206.

Forejt, J., and P. Jansa, 2023 Meiotic Recognition of Evolutionarily Diverged Homologs: Chromosomal Hybrid Sterility Revisited. Mol Biol Evol 40.

Forejt, J., P. Jansa and E. Parvanov, 2021 Hybrid sterility genes in mice (Mus musculus): a peculiar case of PRDM9 incompatibility. Trends Genet 37: 1095–1108.

Fujiwara, K., Y. Kawai, T. Takada, T. Shiroishi, N. Saitou et al., 2022 Insights into Mus musculus Population Structure across Eurasia Revealed by Whole-Genome Analysis. Genome Biol Evol 14.

Gregorova, S., V. Gergelits, I. Chvatalova, T. Bhattacharyya, B. Valiskova et al., 2018 Modulation of Prdm9-controlled meiotic chromosome asynapsis overrides hybrid sterility in mice. Elife 7.

Gupta, S., J. A. Stamatoyannopoulos, T. L. Bailey and W. S. Noble, 2007 Quantifying similarity between motifs. Genome Biol 8: R24.

Hammer, M. F., and L. M. Silver, 1993 Phylogenetic analysis of the alpha-globin pseudogene-4 (Hba-ps4) locus in the house mouse species complex reveals a stepwise evolution of t haplotypes. Molecular Biology and Evolution 10: 971–1001.

Hardouin, E. A., A. Orth, M. Teschke, J. Darvish, D. Tautz et al., 2015 Eurasian house mouse (Mus musculus L.) differentiation at microsatellite loci identifies the Iranian plateau as a phylogeographic hotspot. BMC Evol Biol 15: 26.

Harr, B., E. Karakoc, R. Neme, M. Teschke, C. Pfeifle et al., 2016 Genomic resources for wild populations of the house mouse, Mus musculus and its close relative Mus spretus. Sci Data 3: 160075.

Hayashi, K., K. Yoshida and Y. Matsui, 2005 A histone H3 methyltransferase controls epigenetic events required for meiotic prophase. Nature 438: 374–378.

Herrmann, B. G., and H. Bauer, 2012 The mouse t-haplotype, pp. 297-314 in Evolution of the House Mouse.

Herrmann, B. G., B. Koschorz, K. Wertz, K. J. McLaughlin and A. Kispert, 1999 A protein kinase encoded by the t complex responder gene causes non-mendelian inheritance. Nature 402.

Imai, Y., F. Baudat, M. Taillepierre, M. Stanzione, A. Toth et al., 2017 The PRDM9 KRAB domain is required for meiosis and involved in protein interactions. Chromosoma 126: 681–695.

Jeffreys, A. J., V. E. Cotton, R. Neumann and K. W. Lam, 2013 Recombination regulator PRDM9 influences the instability of its own coding sequence in humans. Proc Natl Acad Sci U S A 110: 600–605.

Jeffreys, A. J., and R. Neumann, 2002 Reciprocal crossover asymmetry and meiotic drive in a human recombination hot spot. Nat Genet 31: 267–271.

Jeffreys, A. J., and R. Neumann, 2005 Factors influencing recombination frequency and distribution in a human meiotic crossover hotspot. Hum Mol Genet 14: 2277–2287.

Jeffreys, A. J., R. Neumann and V. Wilson, 1990 Repeat unit sequence variation in minisatellites: a novel source of DNA polymorphism for studying variation and mutation by single molecule analysis. Cell 60: 473–485.

Kelemen, R. K., M. Elkrewi, A. K. Lindholm and B. Vicoso, 2022 Novel patterns of expression and recruitment of new genes on the t-haplotype, a mouse selfish chromosome. Proc Biol Sci 289: 20211985.

Kelemen, R. K., and B. Vicoso, 2018 Complex History and Differentiation Patterns of the t-Haplotype, a Mouse Meiotic Driver. Genetics 208: 365–375.

Kono, H., M. Tamura, N. Osada, H. Suzuki, K. Abe et al., 2014 Prdm9 polymorphism unveils mouse evolutionary tracks. DNA Res 21: 315–326.

Latrille, T., L. Duret and N. Lartillot, 2017 The Red Queen model of recombination hotspot evolution: a theoretical investigation. Philos Trans R Soc Lond B Biol Sci 372.

Lawal, R. A., U. P. Arora and B. L. Dumont, 2021 Selection shapes the landscape of functional variation in wild house mice. BMC Biol 19: 239.

Lawson, C., M. Gieske, B. Murdoch, P. Ye, Y. Li et al., 2011 Gene expression in the fetal mouse ovary is altered by exposure to low doses of bisphenol A. Biol Reprod 84: 79–86.

Lustyk, D., S. Kinsky, K. K. Ullrich, M. Yancoskie, L. Kasikova et al., 2019 Genomic Structure of Hstx2 Modifier of Prdm9-Dependent Hybrid Male Sterility in Mice. Genetics 213: 1047–1063.

Lyon, M. F., 2003 Transmission ratio distortion in mice. Annu Rev Genet 37: 393–408.

Mihola, O., Z. Trachtulec, C. Vlcek, J. C. Schimenti and J. Forejt, 2009 A mouse speciation gene encodes a meiotic histone H3 methyltransferase. Science 323: 373–375.

Mukaj, A., J. Pialek, V. Fotopulosova, A. P. Morgan, L. Odenthal-Hesse et al., 2020 Prdm9 inter-subspecific interactions in hybrid male sterility of house mouse. Mol Biol Evol.

Muller, H. J., 1942 Isolating Mechanisms, Evolution and Temperature. 71–125.

Myers, S., R. Bowden, A. Tumian, R. E. Bontrop, C. Freeman et al., 2010 Drive against hotspot motifs in primates implicates the PRDM9 gene in meiotic recombination. Science 327: 876–879.

Odenthal-Hesse, L., I. L. Berg, A. Veselis, A. J. Jeffreys and C. A. May, 2014 Transmission distortion affecting human noncrossover but not crossover recombination: a hidden source of meiotic drive. PLoS Genet 10: e1004106.

Olds-Clarke, P., 1997 Models for male infertility: the t haplotypes. Reviews of Reproduction 2: 157–164.

Oliver, P. L., L. Goodstadt, J. J. Bayes, Z. Birtle, K. C. Roach et al., 2009 Accelerated evolution of the Prdm9 speciation gene across diverse metazoan taxa. PLoS Genet 5: e1000753.

Paradis, E., and K. Schliep, 2019 ape 5.0: an environment for modern phylogenetics and evolutionary analyses in R. Bioinformatics 35: 526–528.

Parvanov, E. D., P. M. Petkov and K. Paigen, 2010 Prdm9 controls activation of mammalian recombination hotspots. Science 327: 835.

Parvanov, E. D., H. Tian, T. Billings, R. L. Saxl, C. Spruce et al., 2017 PRDM9 interactions with other proteins provide a link between recombination hotspots and the chromosomal axis in meiosis. Mol Biol Cell 28: 488–499.

Patel, A., X. Zhang, R. M. Blumenthal and X. Cheng, 2017 Structural basis of human PR/SET domain 9 (PRDM9) allele C-specific recognition of its cognate DNA sequence. J Biol Chem 292: 15994–16002.

Persikov, A. V., R. Osada and M. Singh, 2009 Predicting DNA recognition by Cys2His2 zinc finger proteins. Bioinformatics 25: 22–29.

Persikov, A. V., and M. Singh, 2014 *De novo* prediction of DNA-binding specificities for Cys_2_His_2_ zinc finger proteins. Nucleic Acids Res 42: 97–108.

Phifer-Rixey, M., B. Harr and J. Hey, 2020 Further resolution of the house mouse (Mus musculus) phylogeny by integration over isolation-with-migration histories. BMC Evol Biol 20: 120.

Pialek, J., M. Vyskocilova, B. Bimova, D. Havelkova, J. Pialkova et al., 2008 Development of unique house mouse resources suitable for evolutionary studies of speciation. J Hered 99: 34–44.

Planchart, A., Y. You and J. C. Schimenti, 2000 Physical mapping of male fertility and meiotic drive quantitative trait loci in the mouse t complex using chromosome deficiencies. Genetics 155: 803–812.

Powers, N. R., E. D. Parvanov, C. L. Baker, M. Walker, P. M. Petkov et al., 2016 The Meiotic Recombination Activator PRDM9 Trimethylates Both H3K36 and H3K4 at Recombination Hotspots In Vivo. PLoS Genet 12: e1006146.

Quinlan, A. R., and I. M. Hall, 2010 BEDTools: a flexible suite of utilities for comparing genomic features. Bioinformatics 26: 841–842.

Santana-Garcia, W., J. A. Castro-Mondragon, M. Padilla-Galvez, N. T. T. Nguyen, A. Elizondo-Salas et al., 2022 RSAT 2022: regulatory sequence analysis tools. Nucleic Acids Res 50: W670–W676.

Schimenti, J. C., J. L. Reynolds and A. Planchart, 2005 Mutations in Serac1 or Synj2 cause proximal t haplotype-mediated male mouse sterility but not transmission ratio distortion. Proc Natl Acad Sci U S A 102: 3342–3347.

Silver, L. M., 1985 MOUSE t HAPLOTYPES. Annual Review of Genetics.

Smagulova, F., K. Brick, Y. Pu, R. D. Camerini-Otero and G. V. Petukhova, 2016 The evolutionary turnover of recombination hot spots contributes to speciation in mice. Genes Dev 30: 266–280.

Staubach, F., A. Lorenc, P. W. Messer, K. Tang, D. A. Petrov et al., 2012 Genome patterns of selection and introgression of haplotypes in natural populations of the house mouse (Mus musculus). PLoS Genet 8: e1002891.

Tamura, K., and M. Nei, 1993 Estimation of the number of nucleotide substitutions in the control region of mitochondrial DNA in humans and chimpanzees. Mol Biol Evol 10: 512–526.

Trachtulec, Z., C. Vlcek, O. Mihola, S. Gregorova, V. Fotopulosova et al., 2008 Fine haplotype structure of a chromosome 17 region in the laboratory and wild mouse. Genetics 178: 1777–1784.

Turner, L. M., and B. Harr, 2014 Genome-wide mapping in a house mouse hybrid zone reveals hybrid sterility loci and Dobzhansky-Muller interactions. Elife 3.

Turner, L. M., D. J. Schwahn and B. Harr, 2012 Reduced male fertility is common but highly variable in form and severity in a natural house mouse hybrid zone. Evolution 66: 443–458.

Ullrich, K. K., M. Linnenbrink and D. Tautz, 2017 Introgression patterns between house mouse subspecies and species reveal genomic windows of frequent exchange.

Valiskova, B., S. Gregorova, D. Lustyk, P. Simecek, P. Jansa et al., 2022 Genic and chromosomal components of Prdm9-driven hybrid male sterility in mice (Mus musculus). Genetics 222.

Vara, C., L. Capilla, L. Ferretti, A. Ledda, R. A. Sanchez-Guillen et al., 2019 PRDM9 diversity at fine geographical scale reveals contrasting evolutionary patterns and functional constraints in natural populations of house mice. Mol Biol Evol.

Walker, M., T. Billings, C. L. Baker, N. Powers, H. Tian et al., 2015 Affinity-seq detects genome-wide PRDM9 binding sites and reveals the impact of prior chromatin modifications on mammalian recombination hotspot usage. Epigenetics Chromatin 8: 31.

Wang, Y., T. Guo, H. Ke, Q. Zhang, S. Li et al., 2021 Pathogenic variants of meiotic double strand break (DSB) formation genes PRDM9 and ANKRD31 in premature ovarian insufficiency. Genet Med 23: 2309–2315.

Widmayer, S. J., M. A. Handel and D. L. Aylor, 2020 Age and Genetic Background Modify Hybrid Male Sterility in House Mice. Genetics 216: 585–597.

Wooldridge, L. K., and B. L. Dumont, 2022 Rapid evolution of the fine-scale recombination landscape in wild house mouse (Mus musculus) populations. Mol Biol Evol.

Wu, H., N. Mathioudakis, B. Diagouraga, A. Dong, L. Dombrovski et al., 2013 Molecular basis for the regulation of the H3K4 methyltransferase activity of PRDM9. Cell Rep 5: 13–20.

Zelazowski, M. J., and F. Cole, 2016 X marks the spot: PRDM9 rescues hybrid sterility by finding hidden treasure in the genome. Nat Struct Mol Biol 23: 267–269.

